# The GC-content at the 5’ends of human protein-coding genes is undergoing mutational decay

**DOI:** 10.1101/2024.03.12.584636

**Authors:** Yi Qiu, Yoon Mo Kang, Christopher Korfmann, Fanny Pouyet, Andrew Eckford, Alexander F. Palazzo

## Abstract

In vertebrates, most protein-coding genes have a peak of GC-content near their 5’ transcriptional start site (TSS). This feature promotes both the efficient nuclear export and translation of mRNAs. Despite the importance of GC-content for RNA metabolism, its general features, origin, and maintenance remain mysterious. We investigated the evolutionary forces shaping GC-content at the transcriptional start site (TSS) of genes through both comparative genomic analysis of nucleotide substitution rates between different species and by examining human *de novo* mutations. Our data suggests that GC-peaks at TSSs were present in the last vertebrate common ancestor and are largely dictated by recombination patterns. We observe that in primates and rodents, where recombination is directed away from TSSs by PRDM9, GC-content at protein-coding gene TSSs is currently undergoing mutational decay. In canids, which lack PRDM9 and perform recombination at TSSs, GC-content at protein-coding gene TSSs is increasing. These patterns extend into the open reading frame affecting protein-coding regions, and we show that changes in GC-content due to recombination affect synonymous codon position choices at the start of the open reading frame. Our results indicate that although high GC-content in protein-coding genes may be shaped by selective pressures to enhance expression, the dynamics of GC-content in mammals are largely shaped by patterns of recombination.

## Introduction

One of the most striking features of human protein coding genes is the pattern of GC-content present along its length (Palazzo et al., 2024; Palazzo and Kang, 2021). In particular, it has been observed that GC-content is highest at the 5’ end of genes, and that this decreases as one travels downstream and is lowest at the 3’ end of genes (Xia et al., 2003; Louie et al., 2003; Zhang et al., 2004; Kalari et al., 2006; Haerty and Ponting, 2015; Zhu et al., 2009; Palazzo and Kang, 2021; Palazzo et al., 2024). In addition, it has been observed that GC-content is higher in exons than in introns, with it being highest in the first exon and decreasing with every subsequent exon (Palazzo et al., 2024; Zhu et al., 2009). A similar pattern has been observed in introns (Palazzo et al., 2024; Zhu et al., 2009).

GC-content appears to play several different roles in the gene expression pathway. GC-rich promoters, often called CpG islands, have been shown to activate transcription (Fenouil et al., 2012). Moreover, differences between GC-content in exons and introns enhance splicing (Amit et al., 2012). Finally, high GC-content, when present at the 5’ end of intron-poor mRNAs, promote mRNA nuclear export (Palazzo and Akef, 2012; Mordstein et al., 2020; Palazzo and Kang, 2021). Indeed, it has been noted that RNA elements that are GC-rich tend to promote nuclear export when inserted at the 5’ends of intronless reporter mRNAs (Palazzo et al., 2007; Lei et al., 2011, 2013; Tarnawsky and Palazzo, 2012). These GC-rich regions likely recruit protein factors, such as THO complex components, SR proteins and RBM33, which directly recruit nuclear transport receptors, like NXF1/NXT1, that ferry the mRNAs across the nuclear pore (Huang and Steitz, 2001; Huang et al., 2003; Thomas et al., 2022; Xie et al., 2023). This GC-dependent nuclear export is likely also important for the export of certain long non-coding RNAs, such as *NORAD*, which are produced from intronless genes (Zuckerman et al., 2020; Thomas et al., 2022). In contrast, mRNAs from intron-rich genes largely acquire nuclear export factors during splicing (Masuda et al., 2005; Zuckerman et al., 2020). As a result highly spliced mRNAs are exported in a GC-content-independent manner (Mordstein et al., 2020).

The forces that dictate GC-content in genes remain unclear. There is some evidence suggesting that GC-content is partially shaped by adaptive evolutionary forces. This comes from the study of *de novo* genes that are generated when mRNAs are inadvertently copied to cDNA and then reinserted into the genome to form active “retrogenes” (Mordstein et al., 2020). Indeed, it has been observed that these genes have elevated GC-content at their 5’ ends in comparison to their intron-containing counterparts, suggesting that elevation of GC-content can be driven by positive selection to drive their efficient export (Mordstein et al., 2020). Moreover, retrogenes tend to arise from parental genes that have high GC-content at their 5’ends (Kaessmann et al., 2009). Despite this, some observations suggest that GC-content at the start of genes may also be influenced by non-adaptive forces. First, highly spliced mRNAs tend to have high GC-content at their 5’ ends despite the fact that it is not required for export and does not affect expression levels (Mordstein et al., 2020). Second, GC-content extends for quite some distance upstream from the transcriptional start site beyond the promoter region (Louie et al., 2003; Zhang et al., 2004) and downstream into the first intron (Palazzo et al., 2024; Zhu et al., 2009), which are under little selection. Thus it is likely that GC-content is also shaped by non-adaptive forces.

One of these forces is regional differences in mutation rates along genes. For example, it has been shown that CpG dinucleotides surrounding the TSS of active human genes experience fewer substitutions (Polak and Arndt, 2008), likely due to the fact that they are hypomethylated, and that this protects them from mutational decay due to the enhanced repair of deaminated cytosines compared to 5-methylated cytosines (Bellacosa and Drohat, 2015). As CpGs only account for a small fraction of all GC-content surrounding the TSS, even in CpG-rich promoters (known as CpG-islands), this form of mutational bias does not explain the pattern of GC-content seen at the beginning of most human genes.

Another major non-adaptive evolutionary force that can influence local GC-content is GC-biased gene conversion (gBGC) (Duret and Galtier, 2009). This process occurs due to the formation of heteroduplex regions between maternal and paternal chromosomes during homologous recombination. When the two chromosomes are heterozygous, mismatches will form in the heteroduplex which are typically corrected in favor of Gs and Cs over As and Ts by about 70% (Bill et al., 1998; Williams et al., 2015). As a result, G and C single nucleotide polymorphisms surrounding recombination sites are more likely to be transmitted to the next generation, and spread more rapidly in the population thus resembling positive selection. Importantly, recombination is concentrated at certain “hotspots” which contain nucleotide motifs that are bound by PRDM9, a histone methyltransferase (Baudat et al., 2010; Paigen and Petkov, 2018). PRDM9 in turn recruits the topoisomerase SPO11 to generate double stranded breaks (DSB) that initiate homologous recombination (Paiano et al., 2020). Thus GC-content is expected, and is indeed observed to be higher near recombination hotspots due to gBGC (REF).

There are interesting connections between recombination hotspots and TSSs. PRDM9 promotes the trimethylation of H3K4 and H3K36, with the former but not the later, also being present at TSSs. Deletion of PRDM9 in mice led to a shift in recombination from hotspots to TSSs (Brick et al., 2012; Mihola et al., 2019). And finally, although PRDM9 is found throughout the eukaryotic tree, some lineages, such as canids and birds, have lost this gene, and perform recombination at TSSs (Auton et al., 2013; Singhal et al., 2015).

PRDM9 and recombination hotspots also experience accelerated rates of evolution due to the rapid elimination of PRDM9 motifs by DSB-driven biased gene conversion (dBGC) (Myers et al., 2010; Lesecque et al., 2014). Subsequently there is selective pressure on PRDM9 to recognize new motifs (Paigen and Petkov, 2018). Thus the effects of recombination, including gBGC, at any one particular loci is short-lived in evolutionary time, but can affect large parts of the genome over extended times as the concentration of recombination events shifts from one loci to the next.

Here we characterize the GC-content surrounding the TSS of protein-coding genes and infer the nucleotide substitution dynamics of these regions. Although high GC-content in protein-coding genes may be shaped in part by selective pressures to promote mRNA nuclear export and translation, our data suggests that this feature is currently under decay in humans and rodents. Our results indicate that the GC-content surrounding the TSS is largely influenced by patterns of recombination.

## RESULTS

### Analysis of the GC-peak in human protein-coding genes

It is known that in human protein-coding genes, GC-content peaks around the transcription start site (TSS) and slopes down into both the upstream intergenic region and downstream into the first exon and even the first intron (Xia et al., 2003; Louie et al., 2003; Zhang et al., 2004; Kalari et al., 2006; Haerty and Ponting, 2015; Zhu et al., 2009; Palazzo and Kang, 2021; Palazzo et al., 2024). We wanted to examine this GC-peak more closely by metagene analysis and compare it to other gene landmarks. As reported elsewhere, GC-content is highest just downstream of the TSS and descends more-or-less symmetrically on both sides until it plateaus off at 45% (Figure 1A-B). Note that the genome average GC-content is roughly 41%, but that genes tend to be enriched in GC-rich isochores (genomic regions elevated in GC-content) (Bernardi, 2000).

**Figure 1.**
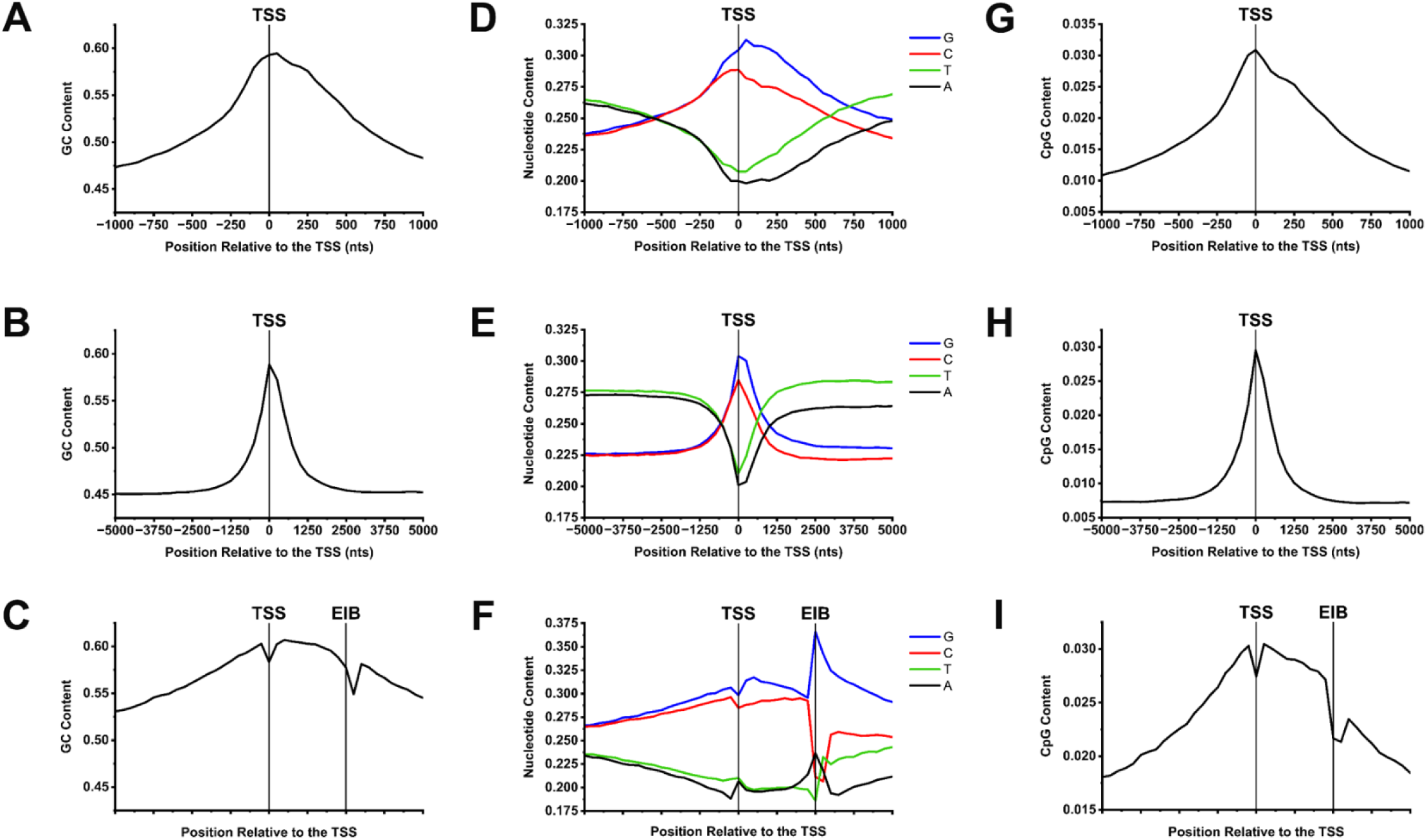
Nucleotide content of genomic regions surrounding TSS sites of human protein-coding genes. For all annotated human protein coding genes (N=18874) GC-content (A-C), nucleotide content on the coding strand (D-F) and CpG-content (G-I) were plotted (*y-axis*) against the nucleotide position (*x-axis*, A-B, D-E, G-H) or along genomic sequence, normalized in length to align all TSSs and first exon-intron boundaries (EIB; *x-axis* C, F, I). Note that for C, F and I, each graph consists of 41 bins – bin size is adjusted for each gene so that the exon occupies 10 bins. The 41 bins consist of 20 upstream, one at the TSS and 20 downstream (10 exonic and 10 intronic).

To get a sense of how GC-content varied with respect to the nearby landmarks, we repeated the analysis of human protein-coding genes but this time we normalized each region to the length of the first exon. We also analyzed an equal amount of sequence into the first intron to examine the GC-content surrounding the exon-intron boundary (EIB). In addition we extended the analysis upstream from the TSS to examine GC-content at the intergenic-exon boundary. Again, we observed that GC-content peaks within the first exon (Figure 1C). Notably, the GC-peak forms a nearly normal curve where it slopes down into both the upstream intergenic region and downstream into the first intron with near identical slopes. This curve is interrupted by the TSS and the exon-intron boundary which are marked by features that are both depressed in GC-content. The dip at the TSS is likely due to the fact that transcription of most genes begins with purines (Carninci et al., 2006; Tamarkin-Ben-Harush et al., 2017) and this slightly elevates the number of Adenines in this region. The dip associated with the exon-intron boundary is likely due to the presence of the 5’ splice site motif, which is slightly GC-poor (Roca et al., 2005).

Next, we examined the content of all four nucleotides over these regions. In particular we wanted to assess any strand asymmetry which manifests as a difference between cognate nucleotides (for example G and C) in each strand. As reported elsewhere, cognate nucleotides were mostly similar upstream of the TSS (Figure 1D) and leveled off at long distances (>2.5kbps) (Figure 1E). Interestingly, divergence of cognate nucleotides was apparent just upstream of the TSS (within 120bps) and this divergence increased downstream of the TSS until it also remained at a certain level for long distances (>2.5kbps) (Figure 1D-E). Since the average exon is only ∼200bps while introns are about an order of magnitude longer, most of the observed strand asymmetries >200bps after the TSS are due to the nucleotide content of the introns. Because these are not under strong selection pressures, the observed asymmetry must be due to strong strand-specific mutation/repair biases that counters mutational decay, which would normally reverse these asymmetries. These biases include transcription-induced damage and/or transcription-coupled DNA repair of the template strand, and enhanced chemical modification of the coding strand, which is single stranded during transcription. Similar arguments have been advanced by other groups (Polak and Arndt, 2008).

When nucleotide composition was examined with respect to the first exon and its delimiting landmarks, further patterns could be discerned (Figure 1F). Right at the TSS, A-content slightly increases likely due to the need for a purine at the first position of the transcript. Within the first exon, A and T displayed little strand asymmetry while G and C diverge – this asymmetry peaks just after the transcriptional start site and then diminishes towards the end of the exon. At the exon-intron boundary, the content of all four nucleotides drastically changes, likely due to the presence of the 5’ splice site motif. Finally, within the intron, strand asymmetries start off very high and then trend towards the strand asymmetries seen in the body of the intron (as seen in Figure 1E).

Next, we examined the level of CpG dinucleotides whose cytosines are often methylated to form 5-methylcytosine. When methylated, CpG are prone to mutational decay as the spontaneous deamination of 5-methylcytosine forms T (Walsh and Xu, 2006). Importantly, CpGs tend to be demethylated around the TSS of active genes, thus reducing their mutational decay. As described previously (Polak and Arndt, 2008), CpG dinucleotides are enriched surrounding the TSS and generally mirror the overall GC-content (Figure 1G-I). Note that CpG dinucleotides make up at most 3% of all sequences around the TSS and thus cannot fully explain the presence of a GC-peak.

### GC-peaks at the 5’ end of genes are present in most vertebrates

To learn how the GC-content of mRNAs varies across vertebrates, we performed a metagene analysis of GC-content along the length of mRNAs from all protein-coding genes from several genomes and compared this to human mRNAs (Figure 2A, for comparison human plots are in gray, species-specific plots are in red). In all species, GC-content in mRNAs was much higher than the genome average (Figure 2A, dotted lines). We calculated the differences between GC content at the beginning of the mRNA compared to the middle of the mRNA for all genes analyzed, and tested whether the distribution of these differences is significantly higher than 0 (Figure 2B). We observed that in amniotes, GC-content was significantly higher at the 5’ end of the mRNA, and the GC-levels gradually decreased downstream (Figure 2A-B). We also observed that this GC-peak was missing in several non-amniotes, including toads, coelacanth, and zebrafish (Figure 2A-B). Since it is present in non-boney fish (e.g. shark and lamprey), it is likely that the GC-peak was present in the vertebrate common ancestor and lost several times (e.g. toads, coelcanths and zebrafish, see asterisks). All species, with the possible exception of turtles and lamprey, had a dip in GC-content at the 3’ end of their mRNAs (Figure 2A).

**Figure 2.**
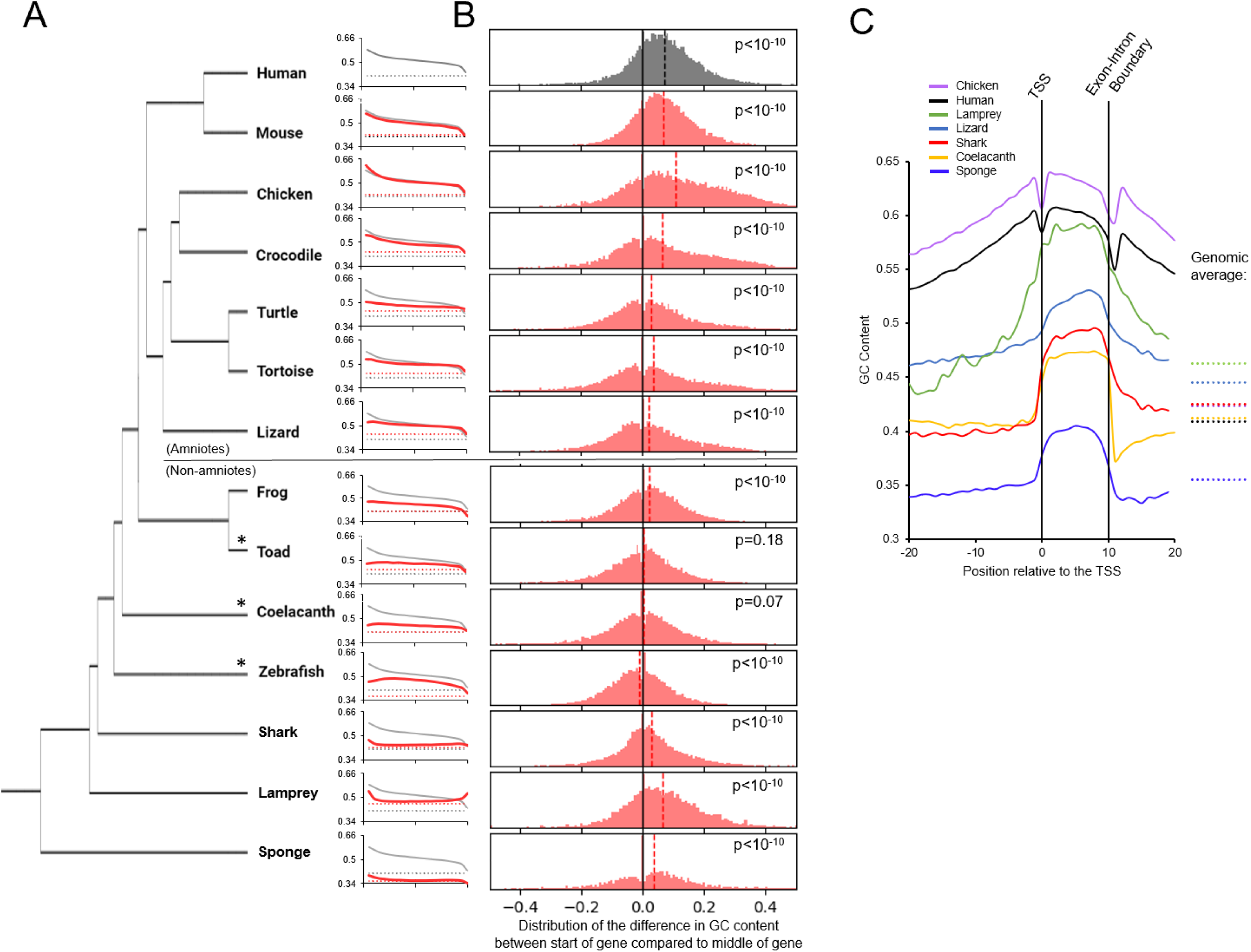
Analysis of GC-content of mRNAs and TSSs from various vertebrate genomes. A) For all annotated mRNAs of the indicated species (red solid lines), the average GC-content (y-axis) was plotted against the normalized mRNA length (from 5’ end to 3’ end in 20 bins; x-axis). Genome average GC-content was also plotted (red dashed lines). As a comparison, GC-content for human mRNA (black solid lines) and human genome (black dashed lines) were replotted in each graph. Phylogenetic relationships of the organisms are indicated on the left B) For each indicated species, the bar graph displays the fraction of mRNAs (y-axis) with various differences in GC-content between the start (bin 1) and the middle (bin 11) of the mRNA (total bins 20; x-axis). Average difference in GC-content for all mRNAs in a given species is indicated by the dashed line. Wilcoxon signed-rank test was performed for the GC content at the start of the mRNA and the middle of the mRNA, and p-values are indicated for each species analyzed. C) For all annotated protein-coding genes of the indicated species, a metaplot of the average GC-content (y-axis) along genomic sequence, normalized in length to align all TSSs and first exon-intron boundaries (as described in Figure 1C; x-axis). As a comparison, genome average GC-content was also plotted (dashed lines on the right).

We next examined GC-content surrounding the first exon and its associated landmarks to examine the GC-peak more closely in a select number of species (Figure 2C). For both humans and chicken, GC-content peaks within the first exon and forms a nearly normal curve where it slopes down into both the upstream intergenic region and downstream into the first intron. Despite this, GC-content in exons, introns and upstream intergenic regions were much higher than the genome average (see dashed lines). Interestingly, this pattern is different in the non-amniotes examined, including anole lizard, coelacanth, shark and lamprey. These organisms had clear differences in GC-content between their first exon and surrounding sequences (upstream and intronic sequences), which came close to the overall genomic GC-content. It is thus likely that amniotes experience specific evolutionary forces that elevate GC-content in regions surrounding the TSS, although we can’t rule out the possibility that these are more ancestral and were lost in most other lineages and hard to see in others. Of course, our ability to detect these trends relies heavily on the correct annotation of these genomes as the majority of the GC-peak should be within the 5’UTR. Nevertheless, we can still detect the GC-peak in human mRNAs if we restrict ourselves to the ORF (Figure S1), and this remains true even if we parse out mRNAs that encode a signal sequence, as the signal sequence coding region is known to be GC-rich and found at the 5’ end of the ORF (Palazzo et al., 2007, 2013; Cenik et al., 2011).

### GC-content in genes correlates with recombination

Recombination is known to elevate GC-content through GC-biased gene conversion (gBGC) (Duret and Galtier, 2009). Although gBGC is thought to be widespread in animals (Galtier et al., 2018), it has specific effects that appear to be amniote-specific (Figuet et al., 2015). To examine whether the frequency of recombination correlates with the GC-peak, we analyzed the GC-content of mRNAs from human genes divided into two categories: those that undergo frequent recombination (top 10%) and those that undergo infrequent recombination (bottom 10%) (Pouyet et al., 2017). The GC-content of mRNAs undergoing frequent recombination is significantly higher than the average, all along the length of the mRNA (Figure 3A), including at the 5’ end (Figure 3A-C) and at the midpoint (Figure 3A, C). The opposite is true for mRNAs undergoing infrequent recombination (Figure 3A-C). The same trend is observed when examining these genes around their TSS, however all genes, whether or not recombination is frequent, have a GC-peak at the 5’ end that is significantly higher in GC-content than the middle of the gene (Figure 3D). These results suggest that recombination affects GC-content throughout the entire length of the gene and not just at the TSS. However since genes that undergo the least amount of recombination have the highest relative GC-peak (GC-content at the 5’ end compared to the middle of the gene), it is unlikely that current recombination patterns are responsible for the presence of the peak.

**Figure 3.**
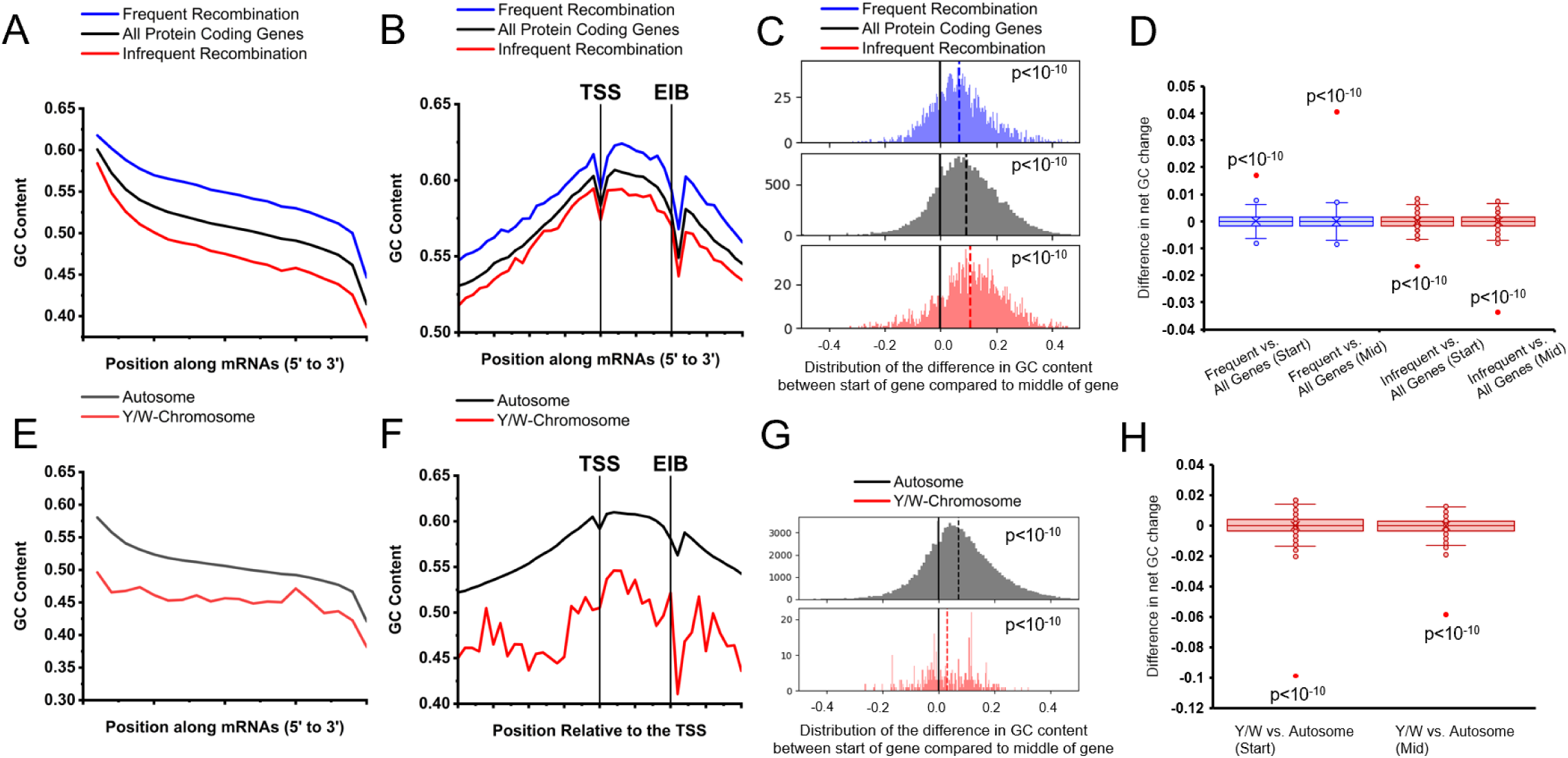
Effects of recombination on the GC-content of mRNAs and TSSs. A) For each set of annotated human mRNAs, the average GC-content (*y-axis*) was plotted against the normalized mRNA length (from 5’ end to 3’ end in 20 bins; *x-axis*). mRNA transcribed from human genes with the 10% highest (blue line) and 10% lowest (red line) recombination frequencies were plotted. The average of all mRNAs was also plotted (black line). B) For each set of annotated human protein-coding genes described in A, the average GC-content (*y-axis*) was plotted against genomic sequence normalized to the TSS and first exon-intron boundary (EIB) over 41 bins (as described in Figure 1C; *x-axis*). C) Wilcoxon signed-rank test was performed for the GC content at the start of the mRNA and the middle of the mRNA described in A, and p-values were indicated. D) Permutation tests between either the start of the mRNA in test groups described in A, or middle of the mRNA in test groups described in A. Distribution of differences from 1000 randomized permutations are displayed in box and whisker plots. Actual differences are plotted with red dots. E) For each set of annotated mRNAs, the average GC-content (*y-axis*) was plotted against the normalized mRNA length of (from 5’ end to 3’ end in 20 bins; *x-axis*). mRNAs transcribed from the autosomes of 5 species (human, mouse, rat, pig, chicken; black line) was compared to mRNAs transcribed from the Y (human, mouse, rat, pig) or W (chicken) chromosomes. F) For each set of annotated protein-coding genes described in E, the average GC-content (*y-axis*) was plotted against genomic sequence normalized to the TSS and first exon-intron boundary (EIB) over 41 bins (as described in Figure 1C; *x-axis*). G) Wilcoxon signed-rank test was performed for the GC content at the start of the mRNA and the middle of the mRNA described in E, and p-values were indicated. H) Permutation tests between either the start of the mRNA in test groups described in E, or middle of the mRNA in test groups described in E. Distribution of differences from 1000 randomized permutations are displayed in box and whisker plots. Actual differences are plotted with red dots.

To further explore this relationship, we next examined protein-coding mRNAs from the Y chromosome, which is not subjected to recombination events. Since there are few genes on the human Y, we compiled genes from Y (mammals) and W (for birds) chromosomes from five species (humans, mice, rat, pigs and chickens). Similar to mRNAs undergoing infrequent recombination, GC-content for Y/W chromosome mRNAs is significantly lower around the TSS and throughout their entire length when compared to those from autosome genes (Figure 3E-H). There is still a significant GC-peak in the Y/W chromosome genes (Figure 3G), however, the GC-peak at the 5’ of the mRNAs is less drastic in Y/W chromosomes compared to autosomes (Figure 3G-H).

Overall, the data are in line with the idea that recombination can affect GC-content, although it does not appear that this explains the GC-peak.

### Primates and rodents are experiencing a loss of GCs surrounding protein-coding gene TSSs

Thus far we have examined the current static picture of GC-content surrounding the TSS. To determine the nucleotide substitution dynamics, we used comparative phylogenetic analysis. We aligned DNA sequences 2500 bp upstream and downstream of the TSS of homologous genes in humans and chimpanzee to identify substitution events. To infer the ancestral state we compared these alignments to the gorilla homolog. Strikingly, we observed that GC-content surrounding the TSS of protein-coding genes is decaying in both humans and chimpanzee (Figure 4A-B). This can be seen even when examining transition substitutions (Figure S2A-B). Note that fluctuations in the total number transition substitutions around the TSS (e.g. A to G and T to C) are due to changes in nucleotide content. When we computed the nucleotide substitution rates by dividing each substitution event over the total mutable nucleotides (e.g. [G to A]/[G]) many of these fluctuations, especially A to G and T to C, are dampened near the TSS (Figure S2G-H), suggesting that the overall loss in GCs was partially due to the presence of fewer mutable A/T nucleotides surrounding the TSS. When random intergenic sequences were analyzed, GC-content was slightly increasing (Figure S3A, B, the averages of these were plotted in Figure 4A-B, dashed lines). We next performed permutation tests between the change of GC at TSSs and intergenic sites that were either GC-matched or not, and observed that the GC-loss at TSSs was significantly greater than at the random intergenic site but not the GC-matched site (Figure S3G). This is consistent with the idea that the GC-peak is experiencing decay at roughly the same rate as intergenic regions with equivalently high GC-content.

**Figure 4.**
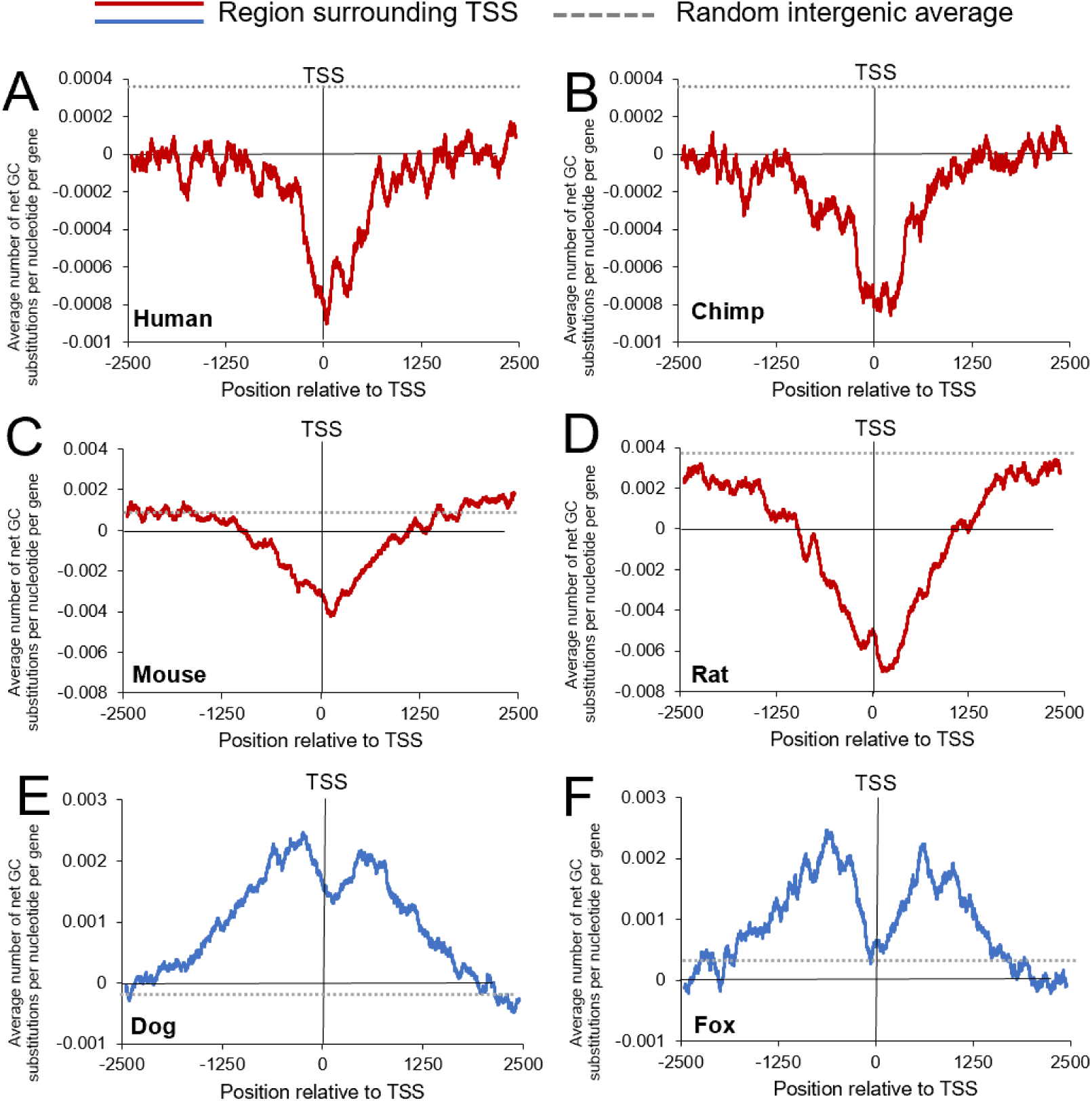
Changes in GC-content surrounding TSSs from various mammalian genomes according to comparative phylogenetic analysis. A-B) Substitution events were inferred by aligning sequences surrounding the TSS of 9248 human and chimpanzee protein-coding genes, using the gorilla sequence to infer the ancestral state. The total change in Gs and Cs in human (A) and chimpanzee (B) were compiled and divided by the number of genes analyzed (*y-axis*) using a sliding window of 100 bp and plotted along genomic regions surrounding the TSS (*x-axis*). The overall average changes in GC-content in random intergenic (dotted line – see Supplemental Figure 3A-B) were plotted. C-D) Similar to (A-B) except that 4840 mouse and rat genes were analyzed using hamster to infer the ancestral state. Random intergenic (dotted line) same as Supplemental Figure 3C-D. E-F) Similar to (A-B) except that 4655 dog and fox genes were analyzed using bear to infer the ancestral state. Random intergenic (dotted line) is the same as Supplemental Figure 3E-F.

To validate these results, we then examined substitutions surrounding the TSSs of mouse and rat genes, using hamster to infer the ancestral state. Again, we observed a decrease in GC-content surrounding the TSS (Figure 4C-D). This can also be seen in the total number of transition mutations (Figure S2C-D). When we computed the nucleotide mutation rates, some of the fluctuations damped, but they were still somewhat pronounced in rat (Figure S2I-J). Again this loss was much higher than at random intergenic sites but not statistically different from GC-matched intergenic sites (Figure S3G). Interestingly, we observe that GC-content is rapidly growing in the genome of rodents, especially in rats (Figure S3C-D).

Although GC-content seemed to decrease at the TSS, it is known to increase near recombination sites due to gBGC (Duret and Galtier, 2009). To confirm this, we examined documented recombination hotspots in the mouse genome (Smagulova et al., 2011), using rat and hamster genomes to infer the substitution dynamics. Indeed, we observed an increase in GC-content in these regions (Figure S4A) that was above the rate for GC-match intergenic regions (Figure S4B, average is replotted as a dashed line in both S4A and S4B).

From these analyses we concluded that GC-content at the TSS of protein-coding genes is not at equilibrium, but in decay in primates and rodents. This decay rate is similar to the decay seen in intergenic regions that have the same GC-content (Figure S3G), this despite the fact that TSSs likely experience fewer CpG deamination events (Polak and Arndt, 2008). Using estimates of divergence between the various organisms, and the number of substitutions, we estimate that this decay in GC-content is on the order of 1% every 10 million years. This implies that the GC-peak built up to some higher point in our evolutionary past, but has since then been in decline.

### *De novo* mutations at human protein-coding gene TSSs show a loss of GC-content

Previously, we examined nucleotide substitutions between related species. To further validate the decay of GC content at the TSS, we mapped human germline *de novo* mutations (DNMs) from 1,548 parents/offspring trios (Jónsson et al., 2017) to regions surrounding protein-coding gene TSSs. In agreement with our substitution analysis, we observed that regions around the TSS had a greater number of mutations that reduced GC-content than gained GC-content (Figure 5A). This difference was significant when compared to random intergenic regions (Figure 5B, Figure 5E solid red bar) but not random GC-matched regions (Figure 5E solid blue bar), suggesting that the peak is undergoing decay at the same rate as regions of the genome that are high in GC-content but not under any selective constraint. This difference could also be seen when total transition mutations were analyzed (Figure S5A TSS, Figure S5C intergenic). We could also see an enhanced number of mutations away from GC at regions further away from the TSS which appeared as shoulders in Figure 5A (nucleotides less than −1250 and greater than 1250). We next computed the nucleotide mutation rates and we observed that transition mutation rates away from G and C were slightly depressed around the TSS (Figure S5B) compared to the rates in intergenic regions (Figure S5D). These rates rose at either end, likely explaining the shoulders in Figure 5A.

**Figure 5.**
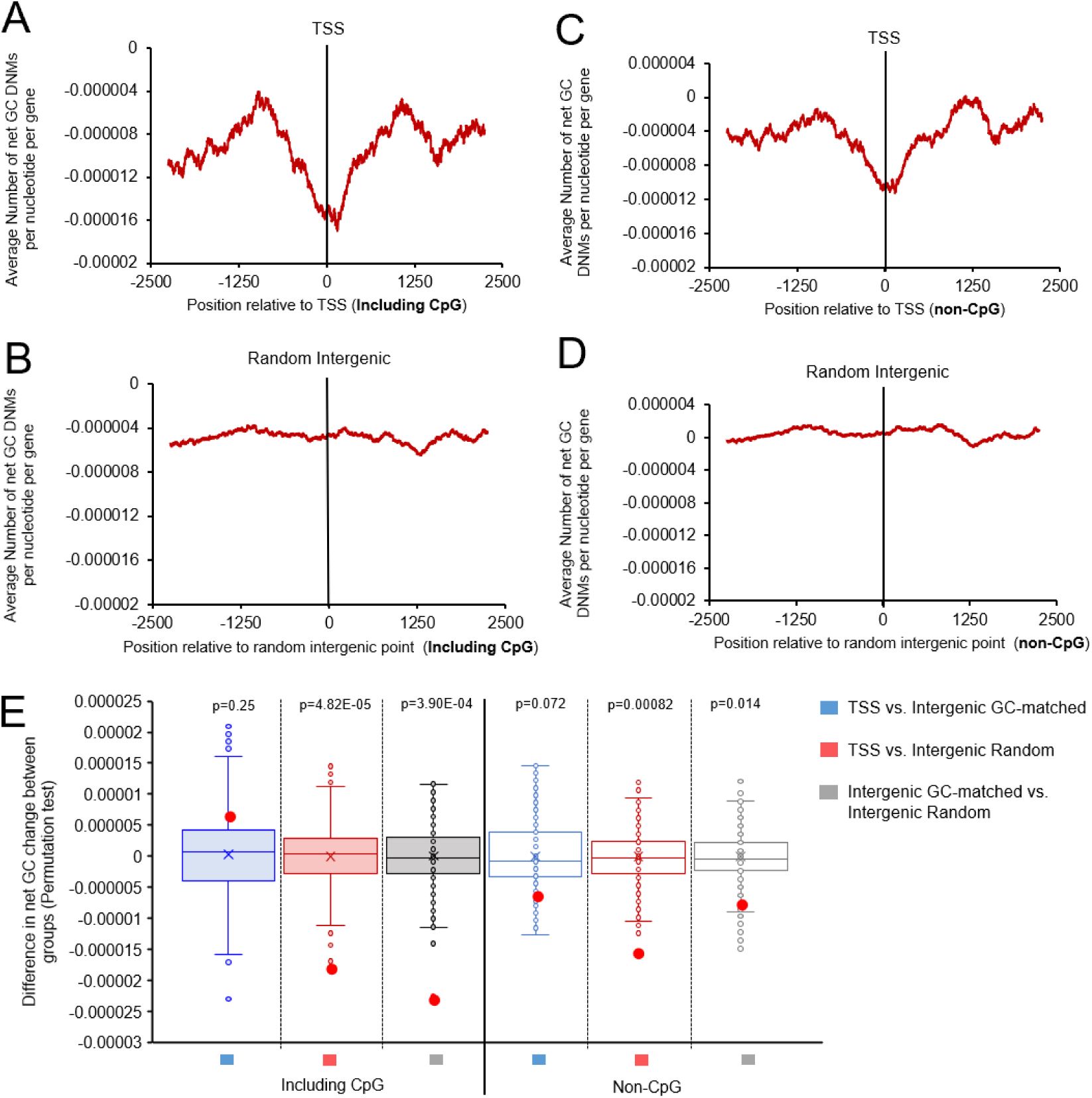
Changes in GC-content due to *de novo* mutations surrounding human TSSs according to parent-offspring trio analyses. A) *De novo* mutations surrounding the TSS of human protein-coding genes were compiled from Jónsson et al., 2017 and divided by the number of genes analyzed (*y-axis*) using a sliding window of 100 bp and plotted along genomic regions surrounding the TSS (*x-axis*). B) Similar to (A) except that random intergenic sequences were analyzed. C-D) Similar to (A-B) except that C to T and G to A mutations in CpGs were omitted. E) Permutation test results comparing net change of GC substitutions around the 0 point for each treatment group in A-D. Distribution of differences from 1000 randomized permutations are displayed in box and whisker plots. Actual differences are plotted with red dots. P-values are indicated.

It was previously observed that CpG decay due to spontaneous deamination was suppressed around TSSs (Polak and Arndt, 2008), likely due to CpG hypomethylation, and this could explain the suppression of G to A and C to T that we observe around the TSSs (Figure S5B). To test this we compiled all non-CpG mutations (TSS shown in Figure 5C, Figure S5E,F, intergenic controls shown in Figure 5C, Figure S5G,H) and found that the excess of mutations away from GC around the TSS was still apparent, however the shoulders seen in Figure 5A were less apparent (Figure 5C). This loss in GC for non-CpG mutations was also significant when compared to random intergenic regions (compare Figure 5C to 5D, compare Figure S5E to S5G, also see stats in Figure 5E empty red bar). There is no significant loss of GC at the TSS for non-CpG mutations when compared to a GC-matched set of intergenic regions (Figure 5E empty blue bar). Overall, our analysis of DNMs confirms that human TSSs are losing GC-content due to mutational decay to a lower equilibrium level.

### Canids are experiencing a gain of GCs surrounding protein-coding gene TSSs

We next examined the GC-dynamics of canids. As described above, PRDM9 became a pseudogene in this clade, and as a result recombination occurs at TSSs in these organisms. We compared sequences from the dog and fox genomes, using bear (which is not a canid) to infer the ancestral state. In marked contrast to what we observed in primates and rodents, the regions surrounding canid protein-coding gene TSSs are gaining GC-content (Figure 4E-F). Again, this trend is also apparent in transition substitutions, both in their total numbers (Figure S2E-F) and in their rates (Figure S2K-L), although there appeared to be a depression of A to G and T to C mutations right around the TSS. The suppression of these mutations may explain the overall dip in GC-gain right in the vicinity of canid TSSs, especially in the fox genome. This dip could be due to negative selection eliminating certain mutations from promoters and exons, and this is somewhat supported by decreases in the other two transition mutations (G to A and C to T, Figure S2E-F, K-L). However, it is also possible that these alterations are due to biased mutation/repair events in the vicinity of the TSS. Despite all these trends, the overall rate of GC-increase in these regions was higher than the genome average, as compiled by measuring the change in GC-content in random intergenic regions (Figure S3E-F, the averages were plotted in Figure 4E-F, dashed lines). When the changes were compared to random GC-matched intergenic regions they were still higher, with the possible exception of the regions right around the fox TSSs (Figure S3G, black and green bars). These changes were statistically significant (Figure S3G).

These data further show that recombination leads to a local increase in GC and that the presence or absence of a functional *PRDM9* gene dictates whether regions surrounding protein-coding gene TSSs are gaining or losing GC-content in mammals.

### Biases due to recombination affect codon usage in the open reading frame

Previously we have shown that we can still detect the GC-peak in human mRNAs if we restrict ourselves to the ORF (Figure S1). Thus, we examined whether the biases caused by recombination extend to the open reading frame (ORF) and affect codon choices. Both adaptive and non-adaptive theories, although not mutually exclusive, have been used to explain synonymous codon usage in humans (Pouyet et al., 2017). The first theory proposes that synonymous codon usage co-adapted with the abundance of tRNAs to optimize the efficiency of translation (dos Reis and Wernisch, 2009). The second theory proposes that synonymous codon usage depends on the large-scale fluctuations of GC-content along chromosomes (Pouyet and Gilbert, 2021).

We performed the same trio-species nucleotide substitution analysis for the entire length of the ORF of all analyzed genes normalized to 40 bins. We observed that human, chimp, mouse and rat (species that have PRDM9 and perform recombination at hotspots) have a significantly higher relative loss of GC-content at the beginning in comparison to downstream portions of the ORF (Figure 6A-D). Interestingly, rodents are gaining GC-content throughout the majority of their ORF, which is in line with our previous data suggesting rodents (especially rats) are gaining GC-content throughout their genome (Figure S3C-D). In species that don’t have PRDM9 and perform recombination at the TSS (dog and fox), there is a significant net gain in GC-content at the beginning of the ORF which diminishes as one travels downstream (Figure 6E,F). We then compared changes in GC-content from ORF confined to either the first exon (and thus close to the TSS) or the fourth exon. We limited our analysis to genes whose start codons occur in the first exon, which accounts for approximately 40% of all human protein-coding genes (Cenik et al., 2011). We observed that human, chimp, mouse and rat all have a significantly higher loss of GC-content in coding sequence from the first exon compared to the fourth exon, whereas dog and fox show the opposite trend (Figure S6A, Table 1). These results suggest that biases dues to the presence or lack of recombination near the TSS affect changes in GC-content that extend into the beginning of the coding regions of genes.

**Figure 6.**
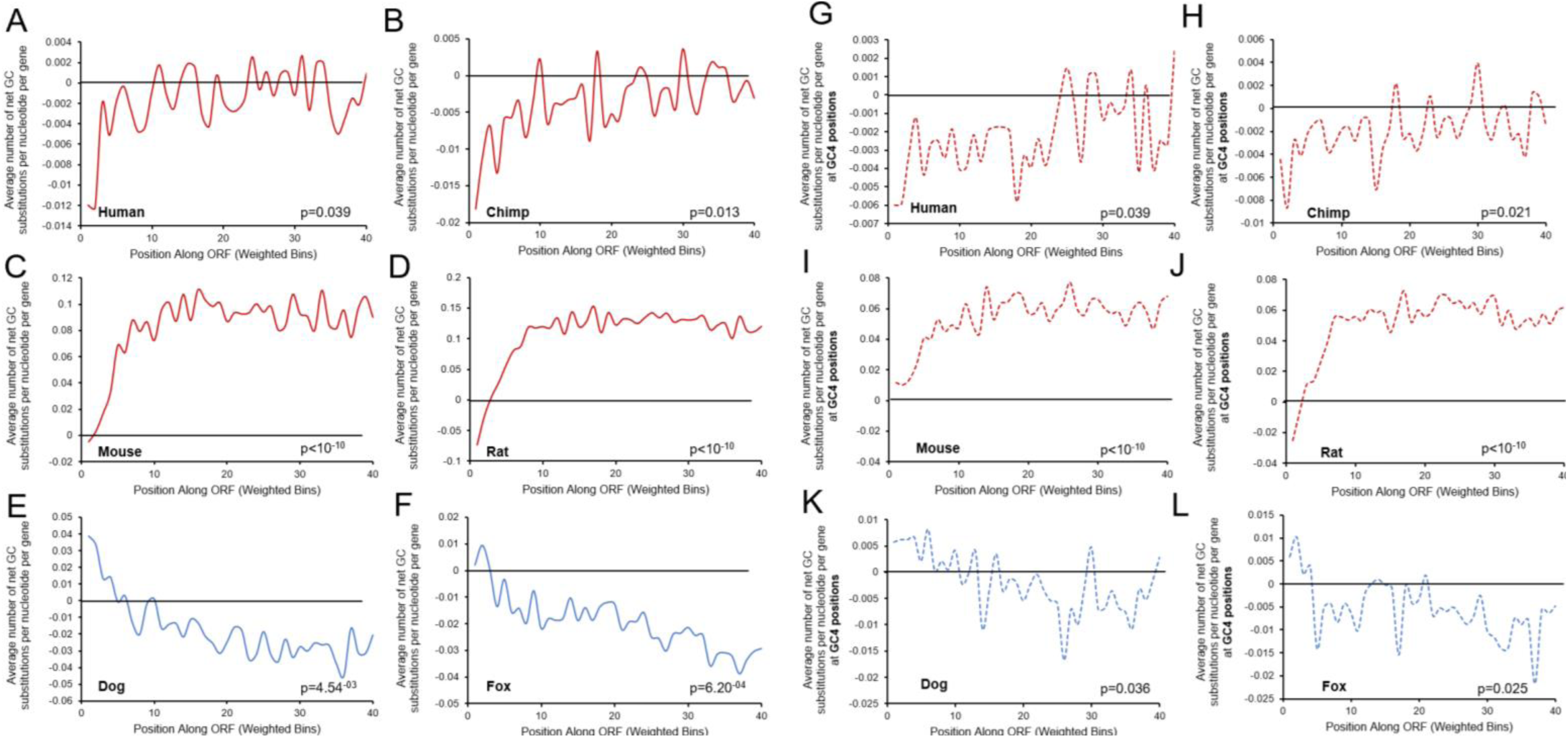
Net change of GC substitutions along the ORF. A-F) Average number of the net change of GC substitutions along the ORF normalized to 40 bins, were plotted for human (A), chimpanzee (B), mouse (C), rat (D), dox (E), and fox (F), with a sliding window of 100 bp. G-L) Average number of the net change of GC substitutions along the ORF at 4-fold degeneracy positions (GC4) normalized to 40 bins were plotted for human (G), chimpanzee (H), mouse (I), rat (J), dox (K), and fox (L), with a sliding window of 100 bp. Only genes with the ORF starting in exon 1 and the start codon (ATG) were included in the analysis. P-values are from permutation tests of changes in GC content between the beginning (bin 1) of the ORF compared to the middle (bin 20) of the ORF (see methods).

**Table 1.** Wilcoxon signed-rank test statistics of changes in GC content between exon 1 and exon 4 of protein coding genes in different organisms according to comparative phylogenetic analyses.

|  | *Human* | *Chimp* | *Mouse* | *Rat* | *Dog* | *Fox* |
| --- | --- | --- | --- | --- | --- | --- |
| *Observed delta (Net change of GC between exon1 and exon 4)* | -0.00024 | -0.00036 | -0.00374 | -0.00662 | 0.000541 | 0.000443 |
| *p-value* | 1.02E-05 | 4.28E-07 | 2.10E-46 | 7.04E-136 | 5.05E-07 | 3.86E-06 |
| *Observed delta (Net change of GC between exon 1 and exon 4 at GC4 positions)* | -0.00028 | -0.00015 | -0.00233 | -0.00312 | 0.000227 | 0.000269 |
| *p-value (GC4)* | 0.005823 | 0.012921 | 4.65E-28 | 1.04E-54 | 0.092441 | 0.037624 |

Next, we examined whether mutational bias affects synonymous codon choice. In particular, we monitored changes in GC-content at 4-fold degeneracy sites of codons (GC4) along the ORF. In agreement with our other findings, we observed that primates and rodents are losing more GCs at GC4 sites in the beginning of the ORF compared to downstream regions (Figure 6G-J), whereas canids show the opposite trend (Figure 6K,L). These observations are all significantly different with the exception of dog, although the same trend is still present (Figure S6B, Table 1). Taken together, we show that changes in GC-content in the coding regions and synonymous codon positions are affected by the presence or lack of recombination at the TSS.

## DISCUSSION

Although there is a tendency in the life sciences to ascribe all features to natural selection, it is clear, but not widely appreciated, that non-adaptive processes also contribute to evolution (Gould and Lewontin, 1979; Lynch, 2007; Koonin, 2016; Palazzo and Kejiou, 2022). In this manuscript we present evidence that the GC-content of protein-coding gene TSSs is influenced, in part, by local rates of recombination. It is possible that local GC-content around the TSS is also subject to selection as high GC-content can promote transcription and mRNA nuclear export. However, it appears that selective forces act in conjunction with (or in some cases in opposition to) non-adaptive processes to shape these GC-peaks.

Our data suggests a model outlined in Figure 7. GC-peaks were likely built up due to evolutionary forces acting on the TSS regions of protein-coding genes in the ancestors of humans and likely all amniotes. It is unlikely that they first arose due to positive selection, which was then relaxed leading to their present day decay. This would require that GC-content play a vital role in the expression of genes in the common ancestor, but not so in organisms such as apes and rodents where these GC-peaks are decaying. Instead, we believe that these peaks were likely caused, and partially maintained, by two main forces. First, CpG that surround the TSS are hypomethylated and thus partially protected from the loss of cytosines by spontaneous deamination. Indeed it has been observed that CpG decay is repressed around TSSs (Polak and Arndt, 2008). Secondly, we believe that in most amniotic lineages, recombination occurred at the TSS of protein-coding genes during distinct time intervals, which were interspaced by other periods where recombination was directed away from TSSs. These distinct periods were likely due to the rapid evolution of PRDM9 (Lesecque et al., 2014). Interestingly, snakes, which have a functional PRDM9, perform recombination at both TSSs and PRDM9-binding sites (Schield et al., 2020). It is believed that there is a tug-of-war competition between TSSs-associated factors and PRDM9 for recruiting downstream factors in the recombination pathway, and that in snakes the strength of these two complexes are somewhat balanced (Hoge et al., 2023). The rapid evolution of PRDM9 may allow different variants, each varying in their relative strength in this tug-of-war competition, to appear in succession within a given lineage. This would cause periodic increases in GC-content at TSSs, in a punctuated manner, followed by intervals of slow decay. Thus, the current alleles of PRDM9 that are present in human, chimp, mouse and rat are very effective at directing recombination away from the TSS to PRDM9-dictated hotspots, and thus these organisms are currently experiencing a decay in their GC-peaks to a lower equilibrium state. Canids, which do not have PRDM9, perform recombination at the TSS, and this further drives up GC-content in these regions to a higher equilibrium state.

**Figure 7.**
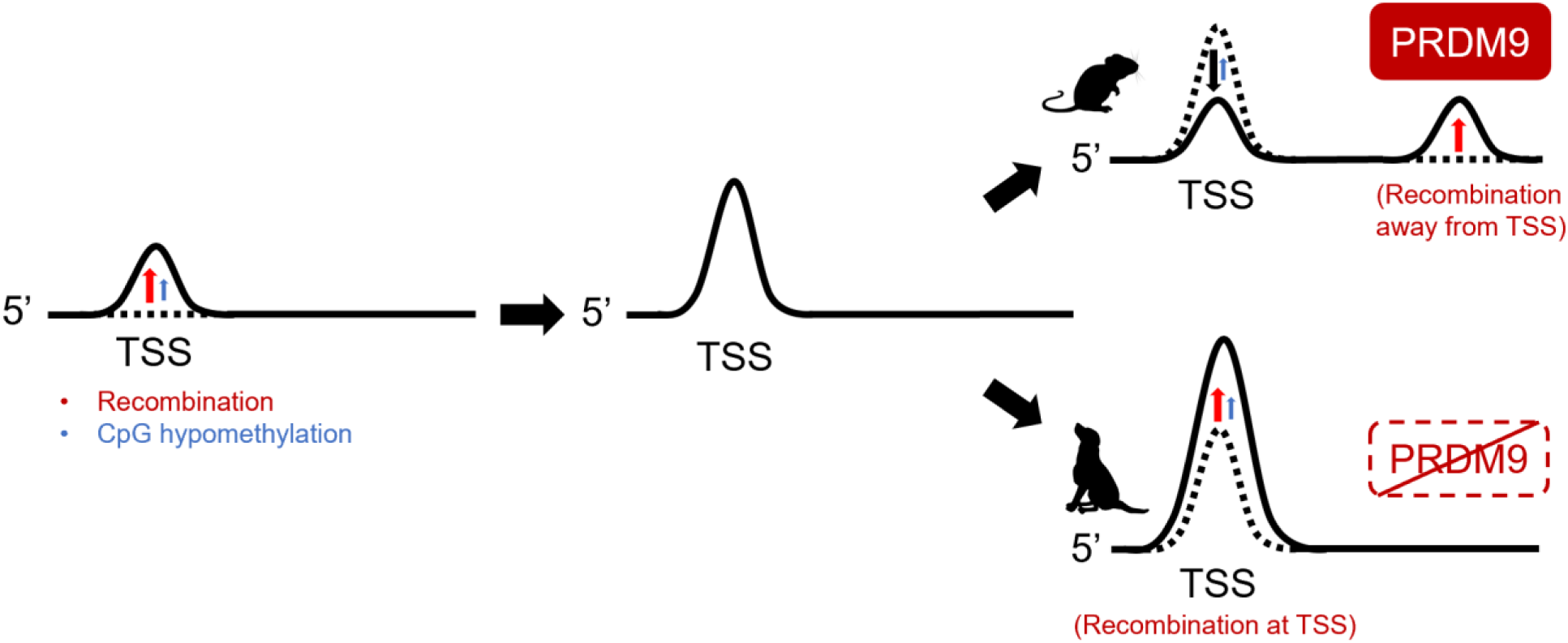
Model for the evolution of GC-Peaks at protein-coding gene TSSs. Recombination shapes GC content at the TSS. The GC peak in the TSS was built up due to recombination and likely to some extent CpG hypomethylation (repression of CpG decay) near promoters. Since then, PRDM9 is directing recombination away from the TSS, leading to a decrease in GC content at the TSS for primates and rodents due to mutational decay (black arrow). In organisms that lost PRDM9, recombination occurs at the TSS, promoting further increase in GC content in canids.

This model is supported by findings in a recent preprint, which documents the equilibrium state of GC-content in TSS regions from numerous organisms. They observe that most mammals have a high GC-content equilibrium state with the exception of humans, mice, some felids and cetaceans. This suggests that most mammals experience some boost in GC-content due to a certain degree of recombination at TSSs despite the fact these organisms contain functional versions of PRDM9 (Joseph et al., 2023).

Finally, our work suggests that certain features of genes, which are recognized by cellular machinery, may be largely shaped by non-adaptive processes. Thus the high GC-content that is found at the start of protein-coding genes, which is used to promote efficient nuclear export of certain mRNAs, appears to be shaped by historical patterns of recombination. It is generally assumed that such “functional” genomic features are at the very least maintained by purging selection. Our new results suggests that some of these features, which appear to specify functional parts of the genome, could be present regardless of selective forces. Instead, parts of the cellular machinery, in this case the mRNA export machinery, seemed to have evolved to recognize these patterns.

It is likely that non-adaptive forces may create other genomic features that are exploited by the cellular machinery for functional ends. These non-adaptive evolutionary forces may help to generate very strong signals in genomes that experience weak selection regimes.

## DATA AVAILABILITY

All python codes can be obtained from the online repository: https://github.com/tinaqiu221/GC_evolution

All raw data can be obtained from the online repository: https://doi.org/10.5281/zenodo.10694966

## METHODS

### Sequence data and annotation

Sequences of animal genomes were accessed and downloaded from the Ensembl database (www.ensembl.org). The genome assembly versions are listed in Table 2. Annotations for protein-coding genes were retrieved using the University of Santa Cruz (UCSC) Genome Browser annotation track database based on the corresponding genome assemblies. For human and mouse, annotations for the TSS were obtained from the FANTOM5 project, which uses Cap Analysis of Gene Expression (CAGE) sequencing (Noguchi et al., 2017). The best TSS from CAGE-seq was defined as the transcript with the highest tags per million score. For all other organisms, annotations for the best TSS were determined by the most commonly used transcript and start site obtained from Ensembl (http://www.ensembl.org/info/data/ftp/index.html/). The exon/intron boundaries are obtained from the corresponding best TSS. Sequences surrounding the TSS were retrieved using the BEDTools suite. (Github: get_homologous_sequences.py) Similarly, intergenic genome regions were retrieved by randomly generating genome coordinates outside of protein-coding regions (Github: generate_random_coordinates.py). Intergenic GC-matched sequences were obtained by selecting from the list of random intergenic sequences, a new set of sequences that have the same distribution of GC-content as the dataset that it is being matched to (Github: gc_match.py). Annotations for the frequency of recombination rate of protein-coding genes were obtained from the HapMap genetic map, and divided into the top 10% and bottom 10% of recombination rates (RRID: SCR_002846) (International HapMap Consortium et al., 2007; Pouyet et al., 2017). Annotations for mouse recombination hotspots were obtained from previously published datasets (Smagulova et al., 2011). Sequences for the open reading frame were obtained using codes from the GitHub repository by Graham E Larue (https://github.com/glarue/cdseq). Genes with their ORF starting in exon 1 were selected by filtering for ORF sequences with start coordinates between the TSS and first exon/intron boundary (ORF_start_exon1.py).

**Table 2.** Genome releases and assembly versions.

| **Organism** | **Genome assembly version** |
| --- | --- |
| Human (*Homo sapien*) | GRCh38 |
| Chimpanzee (*Pan troglodytes*) | Pran_trio3.0 |
| Gorilla (*Gorilla gorilla*) | gorGor4 |
| Mouse (*Mus musculus*) | GRCm39 |
| Rat (*Rattus norvegicus*) | mRatBN7.2 |
| Hamster (*Mesocricetus auratus*) | MesAur1.0 |
| Dog (*Canis lupus familiaris*) | ROS_Cfam_1.0 |
| Fox (*Vulpes vulpes*) | VulVul2.2 |
| Bear (*Ursus americanus*) | ASM334442v1 |
| Chicken (*Gallus gallus*) | bGalGal1.mat.broiler.GRCg7b |
| Crocodile (*Crocodylus porosus*) | CroPor_comp1 |
| Turtle (*Pelodiscus sinensis*) | PelSin_1.0 |
| Tortoise (*Chelonoidis abingdonii*) | ASM359739v1 |
| Lizard (*Podarcis muralis*) | PodMur_1.0 |
| Frog (*Xenopus tropicalis*) | UCB_Xtro_10.0 |
| Toad (*Leptobranchium leishanense*) | ASM966780v1 |
| Coelacanth (*Latimeria chalumnae*) | LatCha1 |
| Zebrafish (*Danio rerio*) | GRCz11 |
| Shark (*Callorhynchus milii*) | Callorhynchus_milii-6.1.3 |
| Lamprey (*Petromyzon marinus*) | Pmarinus_7.0 |
| Sponge (*Amphimedon queenslandica*) | v1.1 |

### Binning strategy and GC content analysis

To assess the GC-content of sequences, we divided the sequence track into different numbers of equally sized bins: 20 bins for mRNA sequences, 41 bins for sequences surrounding the TSS and exon/intron boundary, and 40 bins for ORF sequences. For sequences surrounding the TSS and exon/intron boundary, we calculated the size of the first exon, and obtained size-matched sequences of the corresponding first intron, and two times the size-matched sequences of the corresponding upstream intergenic regions. The 41 bins consist of 20 upstream, one at the TSS and 20 downstream (10 exonic and 10 intronic).We subsequently calculated the GC percent for each bin. (Github: gc_content_binning.py) Whenever multiple isoforms of the same gene are present, the GC content for that gene is the weighted average of all isoforms (e.g. for a gene with 4 isoforms, each isoform contributes to 25% of the GC content for that gene).

### Sequence alignment

Homologous sequences were aligned between primate trios (human, chimpanzee, gorilla), rodent trios (mouse, rat, hamster), and carnivora trios (dog, fox, bear). For sequences surrounding the TSS in protein-coding genes, homologous genes were obtained from Ensembl biomart. For sequences in random intergenic regions and recombination hotspots, triple homologous search was performed using standalone Basic Local Alignment Search Tool (BLAST) from the National Center for Biotechnology Information (NCBI) (Github: get_blast_sequences.py). Triple sequence alignment was performed using the Needleman Alignment algorithm (Github: needleman_alignment.py). A 60% alignment cutoff was applied to all sequence alignments. The total number of trio gene/sequence alignments obtained after the cutoffs used for mapping is in Table 3.

**Table 3.** Number of trio gene/sequence alignments used for mapping.

| **Genome region** | **Organism** | **Number of trio genes/sequences** |
| --- | --- | --- |
| TSS | Human/Chimp | 9248 |
|  | Mouse/Rat | 4840 |
|  | Dog/Fox | 4655 |
| Intergenic random | Human/Chimp | 20209 |
|  | Mouse/Rat | 6142 |
|  | Dog/Fox | 8000 |
| Intergenic GC-matched to TSS | Human/Chimp | 400 |
|  | Mouse/Rat | 509 |
|  | Dog/Fox | 470 |
| Recombination hotspots | Mouse/Rat | 2668 |
| Intergenic GC-matched to recombination hotspots | Mouse/Rat | 4543 |
| ORF | Human/Chimp | 17690 |
|  | Mouse/Rat | 18890 |
|  | Dog/Fox | 15074 |
| ORF (GC4) | Human/Chimp | 14394 |
|  | Mouse/Rat | 8932 |
|  | Dog/Fox | 5876 |
| ORF (Exon 1 and Exon 4) | Human | 5433 |
|  | Chimp | 9240 |
|  | Mouse | 5531 |
|  | Rat | 7113 |
|  | Dog | 6624 |
|  | Fox | 7558 |
| ORF (Exon 1 and Exon 4, GC4) | Human | 4400 |
|  | Chimp | 7648 |
|  | Mouse | 2784 |
|  | Rat | 3912 |
|  | Dog | 2189 |
|  | Fox | 2992 |

### Nucleotide substitution mapping

Gorilla was used as the reference sequence for substitutions in human and chimpanzee, hamster as the reference sequence for substitutions in mouse and rat, and bear used as the reference sequence for dog and fox. Nucleotide substitutions were identified based on the above triple alignments. Transition substitutions, or net change in GC substitutions per gene or sequence analyzed were mapped with a 100 nucleotide sliding window surrounding the TSS, random intergenic site, or recombination hotspot. For rates of substitutions, the number of single substitutions was divided by the total number of that nucleotide of interest per position (eg. number of A to G substitutions for position 1, divided by the total number of A nucleotides in all genes at position 1). Subsequently a 100-nucleotide sliding window is applied. (Github: find_substitutions.py) For the GC4 positions in the ORF, we used only ORF sequences that begin with the start codon “ATG” with sequence lengths divisible by 3. GC substitutions for ORF GC4 positions were extract based substitutions that fall at a position where the sequence is a 3^rd^ synonymous codon position. (Github: find_substitutions_orf_gc4.py). For total substitutions found in exon 1 or exon 4 of the ORF, we used only ORF sequences that contain 4 or more exons. (Github: count_substitutions_per_exon.py).

### De novo mutation mapping

Human germline *de novo* mutations from 1548 parents/offspring trios were obtained from (Jónsson et al., 2017) and mapped to 5kb surrounding the best TSS (defined by FANTOM5 CAGE-seq, see *Sequence data and annotation* section) of protein-coding genes. DNMs were separated into whether or not they occur in a CpG dinucleotide. Rate of DNMs was calculated by dividing the number of DNMs by the number of total mutable nucleotides, similarly to the above nucleotide substitution mapping. (Github: DNM.ipynb).

### Statistical analysis

To statistically test whether the GC content at one locus of a sequence is significantly different from the GC content at another locus, the Wilcoxon signed-rank test is performed between the distributions of the 2 loci (Github: wilcoxontest.py).

To statistically test whether the number of net change in GC substitutions or DNMs at relative nucleotide position 0 is significantly different between two datasets (eg. between the TSS and random intergenic position), a permutation test is performed. In this test, net change in GC substitutions or DNMs from datasets A and B are randomly shuffled into two groups 1000 times, and the differences between the two random groups from the 1000 randomizations are used to generate a normalized curve. The actual observed delta between datasets A and group B is plotted onto the normalized curve, and the p-value is the area under the curve separated by the line x=observed delta. (Github: permutation_test.py) Permutation tests have been described previously. (Wilcox, 2022)

## ACKNOWLEDGEMENTS

We would like to thank Shuyi Shi for help with the *de novo* mutation analysis. This work was funded by a grant to A.F.P. from the Natural Sciences and Engineering Research Council of Canada (FN 492860) and the Jean D’Alembert Fellowship program (France 2030 program ANR-11-IDEX-0003).

## SUPPLEMENTARY FIGURES

**Supplementary Figure 1.**
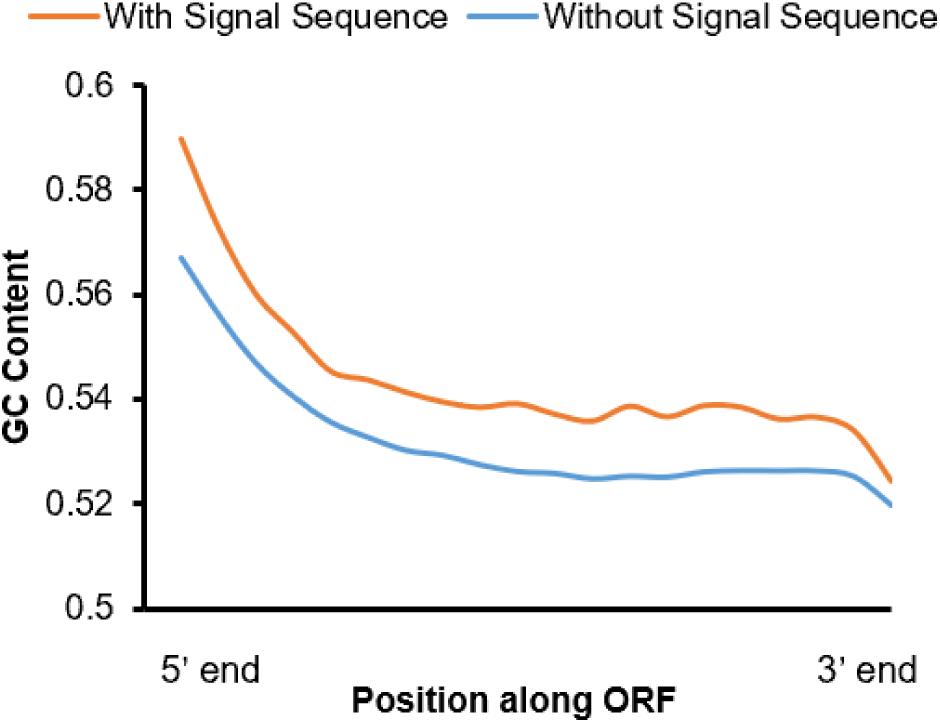
GC content in the open reading frame. The average GC-content for all human protein coding genes (y-axis) was plotted against the normalized open reading frame length (from 5’ end to 3’ end in 20 bins; x-axis), further divided into genes that contain a signal sequence coding region (N=5754) and those without (N=17627).

**Supplementary Figure 2.**
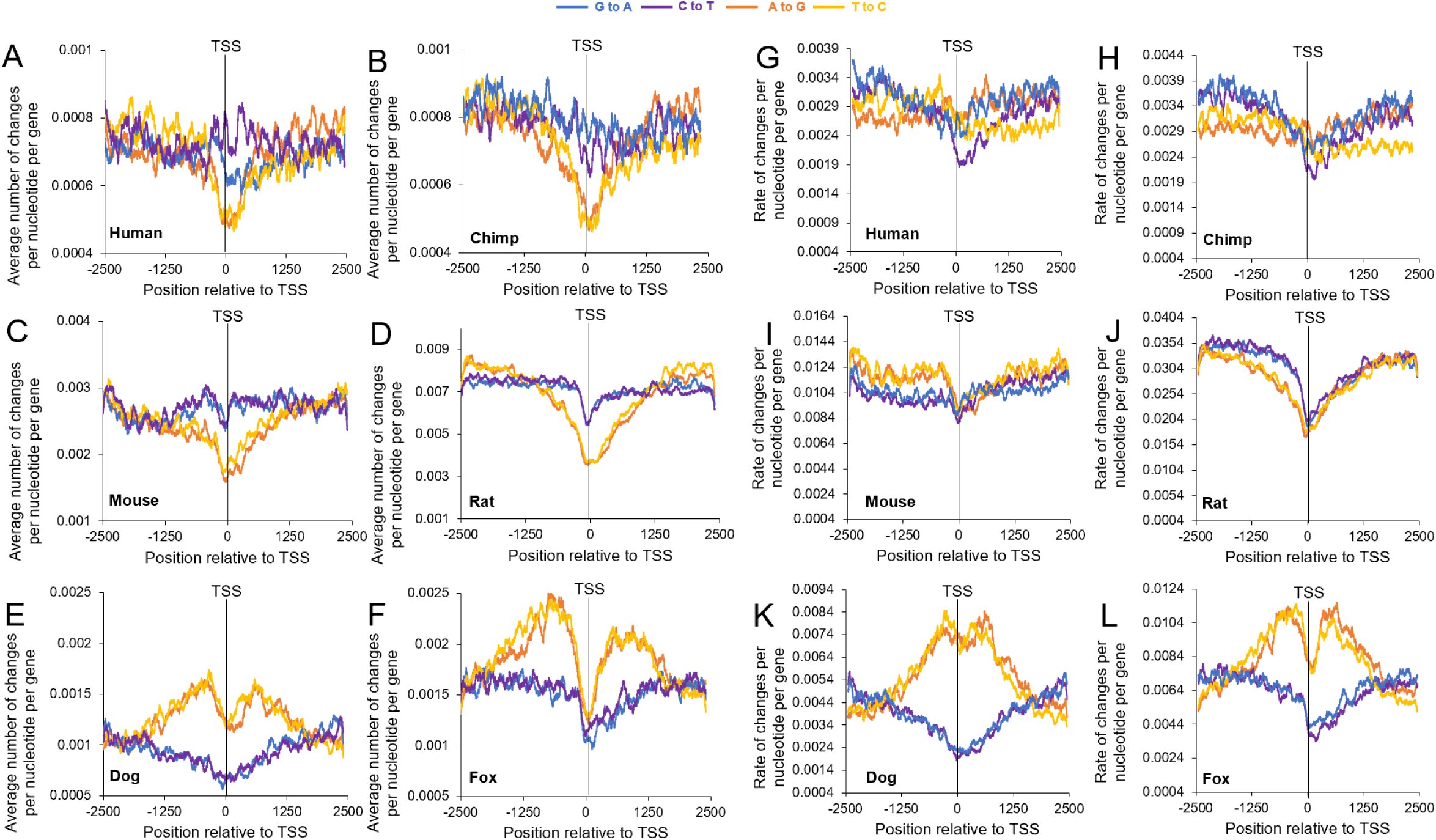
Transition substitution numbers and rates surrounding TSSs from various mammalian genomes according to comparative phylogenetic analyses. A-F) Average number of A to G (orange), T to C (yellow), G to A (blue) and C to T (purple) substitutions in human (A), chimpanzee (B), mouse (C), rat (D), dox (E), and fox (F) divided by the number of genes analyzed (*y-axis*) using a sliding window of 100 bp and plotted along genomic regions surrounding the TSS (*x-axis*). G-L) Rates of each substitution (number of nucleotide substitutions/total mutable nucleotides, e.g., [G to A]/[G]) from A-F were plotted around the TSS for the species indicated.

**Supplementary Figure 3.**
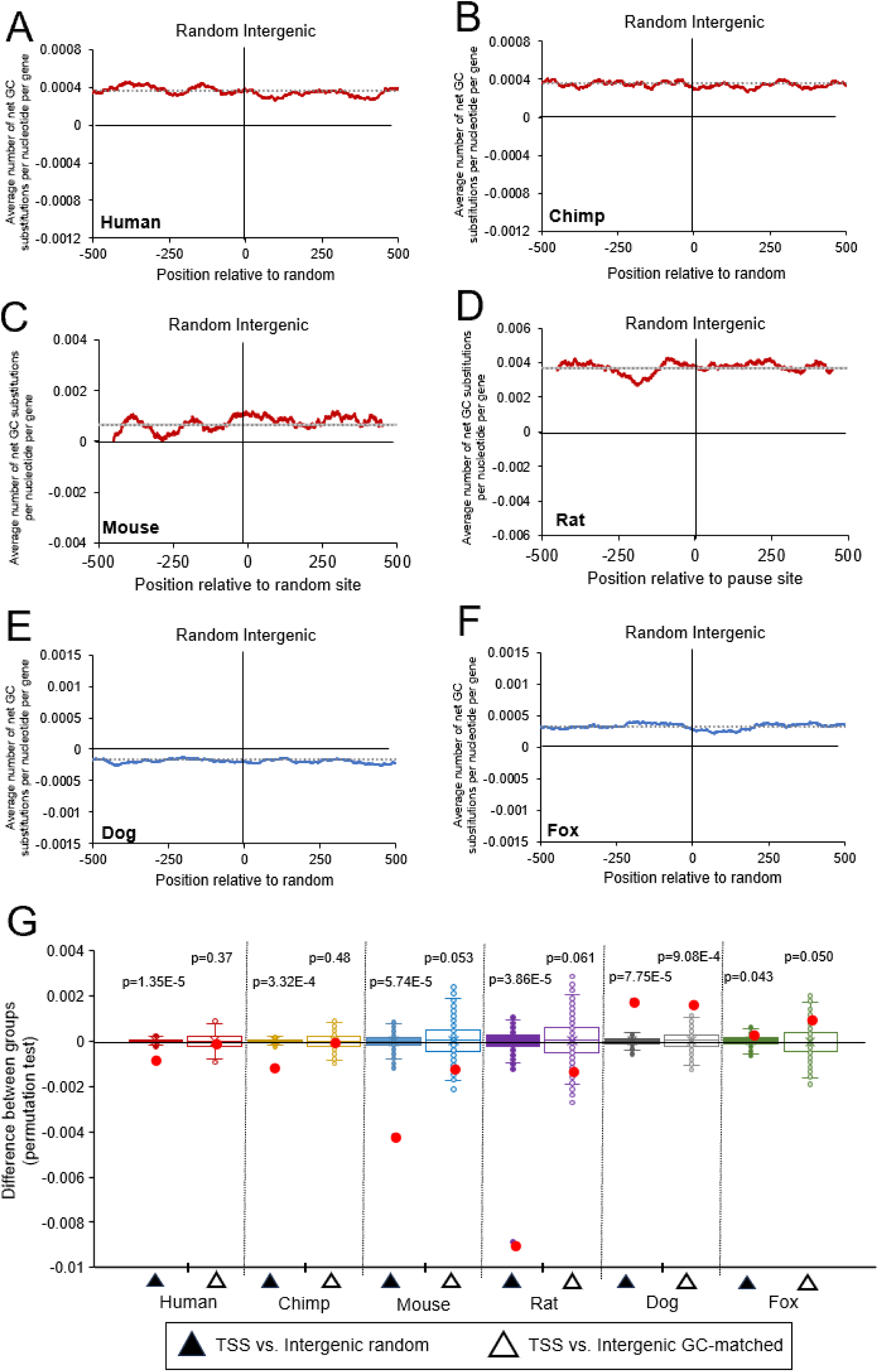
Changes in GC-content surrounding intergenic regions from various mammalian genomes according to comparative phylogenetic analyses. A-F) The total change in Gs and Cs in human (A), chimpanzee (B), mouse (C), rat (D), dox (E), and fox (F) were compiled and divided by the number of genes analyzed (*y-axis*) using a sliding window of 100 bp and plotted along genomic regions surrounding random intergenic points (*x-axis*). Total number of intergenic regions is equal to the number of analyzed genes for each species in Figure 4A-F. The overall average change in GC-content for each plot (dotted line) were plotted – these are also plotted in Figure 4A-F. G) Permutation test results comparing net change of GC substitutions around the 0 point for each treatment group in (A-F). Distribution of differences from 1000 randomized permutations are displayed in box and whisker plots. Actual differences are plotted with red dots. P-values are indicated. Intergenic GC-matched plots are not shown.

**Supplementary Figure 4.**
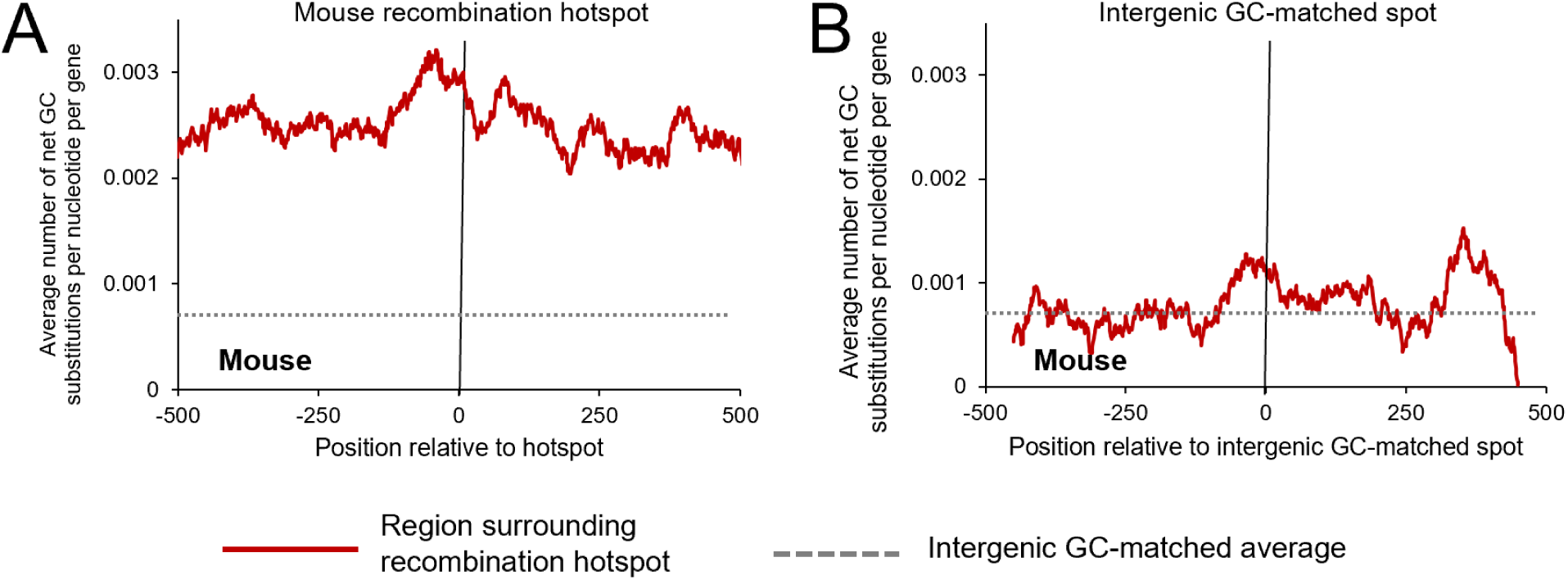
Changes in GC-content surrounding mouse recombination hotspots according to comparative phylogenetic analysis. A) The total change in Gs and Cs in mouse from mouse-rat-hamster trio analysis, were compiled surrounding recombination hotspots (N=2668). Change in G and C divided by the number of regions analyzed was plotted (*y-axis*) over a sliding window of 100 bp along genomic regions surrounding recombination hotspots (*x-axis*). The overall average change in GC-content in mouse at a random intergenic spot (dotted line) was plotted. B) Similar to (A) except plotted along intergenic regions that are GC-matched to the GC content of house recombination hotpots (*x-axis*) (N=4543).

**Supplementary Figure 5.**
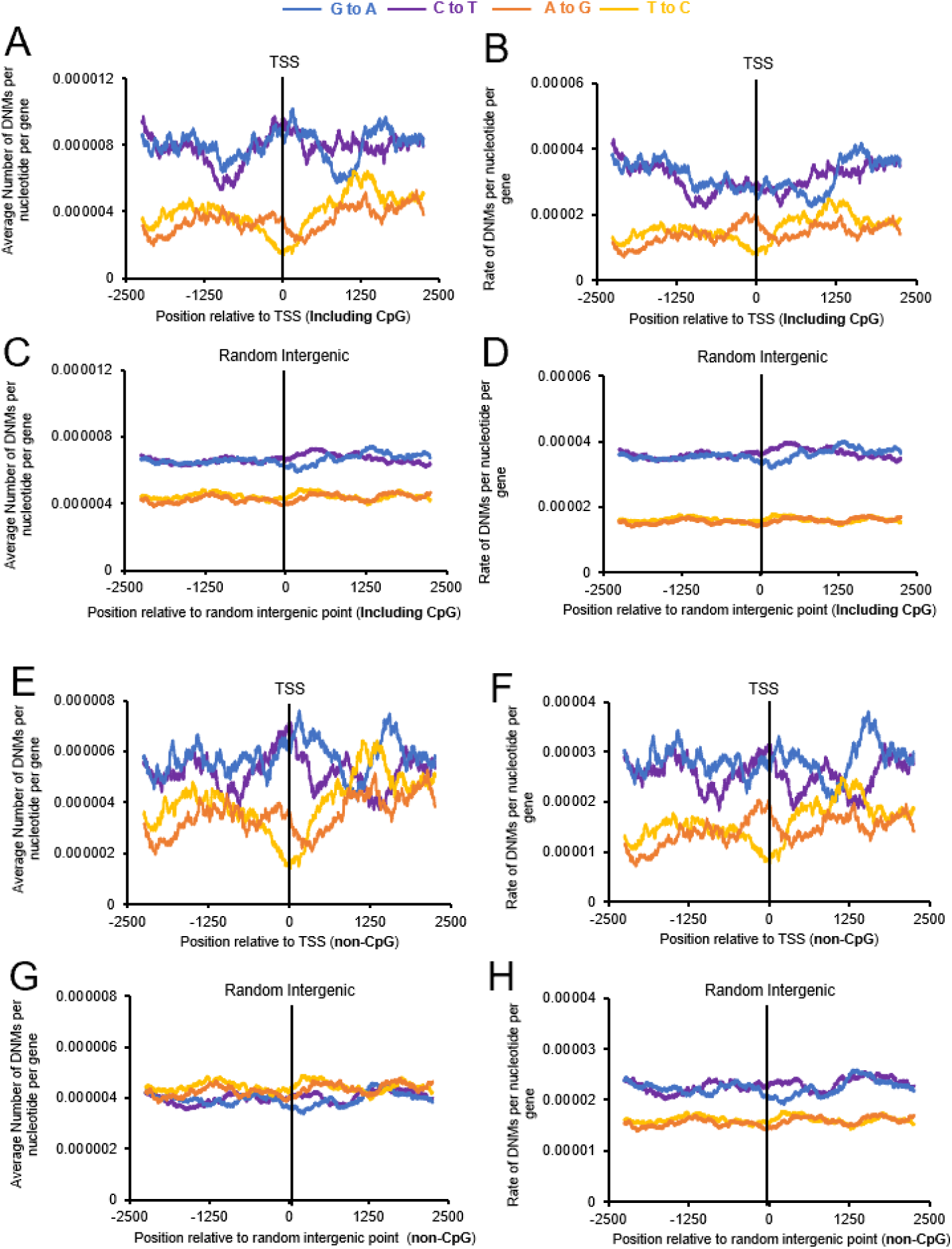
Transition *de novo* mutations, numbers and rates, surrounding human TSSs according to parent-offspring trio analyses. A) Average number of A to G (orange), T to C (yellow), G to A (blue) and C to T (purple) human *de novo* mutations divided by the number of genes analyzed (*y-axis*) using a sliding window of 100 bp and plotted along genomic regions surrounding the TSS (*x-axis*). B) Rates of each *de novo* mutation (number of nucleotide substitutions/total mutable nucleotides, e.g., [G to A]/[G]) from (A). C-D) Similar to (A-B) except plotted around random intergenic region (*x-axis*). E-H) Similar to (A-D) except that C to T and G to A mutations in CpGs were omitted.

**Supplementary Figure 6.**
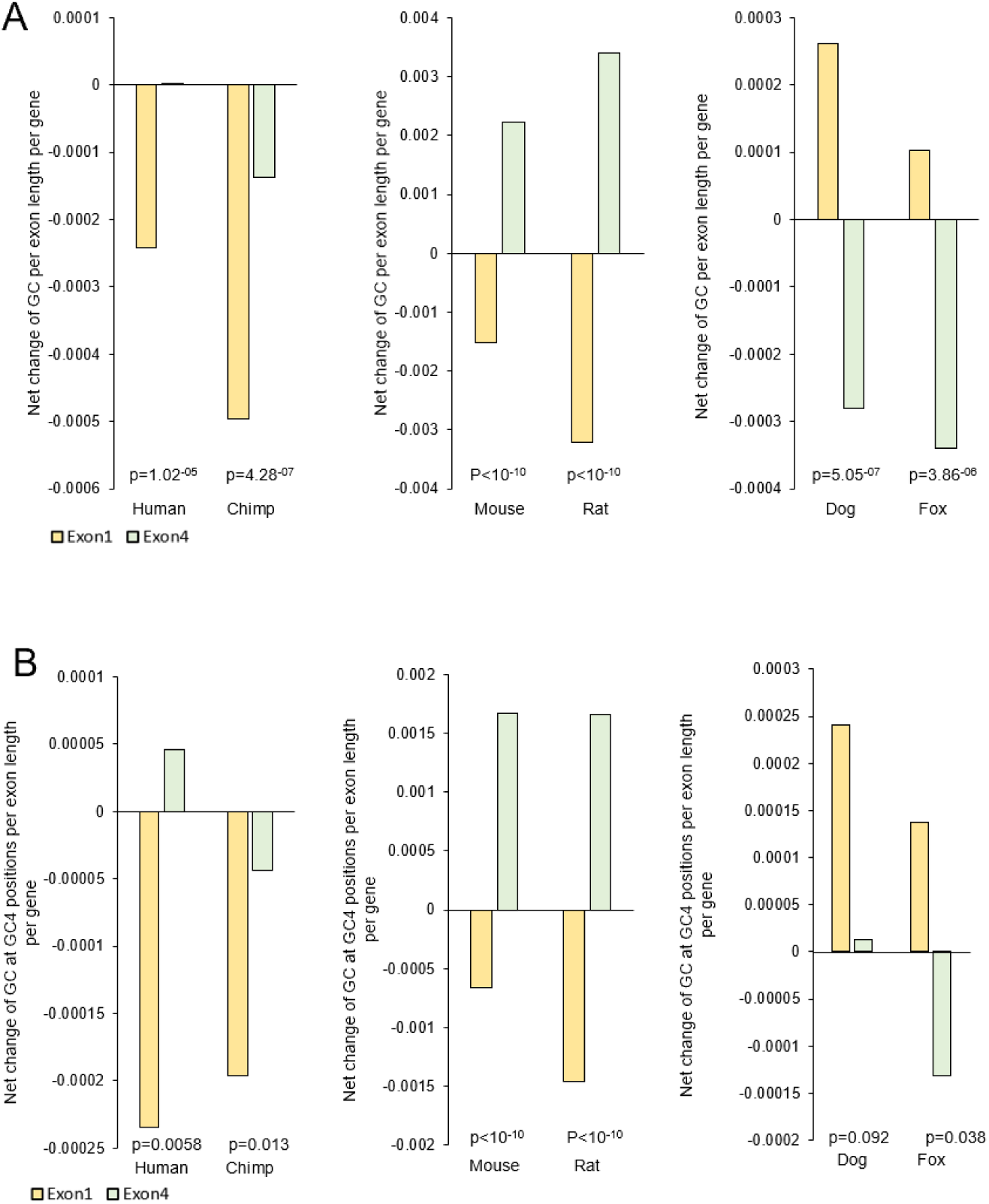
Change in GC content in exon 1 and exon 4 of protein coding genes in different organisms according to comparative phylogenetic analyses. A) The total change in Gs and Cs in exon 1 and exon 4 of genes, divided by the lengths of the corresponding exons and the total number of genes analyzed (Table 3). Only genes with the ORF starting in exon 1, and genes that contain at least 4 exons, were included in the analysis. B) Similar to (A) except only analyzing 4-fold degenerate codon positions (GC4) in exon 1 and exon 4 (Table 3). P-values are from Wilcoxon signed-rank tests of changes in GC content between exon 1 and exon 4 of protein coding genes in different organisms (see methods, Table 1).

## Notes

https://doi.org/10.5281/zenodo.10608324

https://github.com/tinaqiu221/GC_evolution

### Competing Interest Statement

The authors have declared no competing interest.

## REFERENCES

Amit M, Donyo M, Hollander D, Goren A, Kim E, Gelfman S, Lev-Maor G, Burstein D, Schwartz S, Postolsky B, Pupko T, Ast G. 2012. Differential GC Content between Exons and Introns Establishes Distinct Strategies of Splice-Site Recognition. Cell Reports 1:543–556. doi:10.1016/j.celrep.2012.03.013

Auton A, Rui Li Y, Kidd J, Oliveira K, Nadel J, Holloway JK, Hayward JJ, Cohen PE, Greally JM, Wang J, Bustamante CD, Boyko AR. 2013. Genetic recombination is targeted towards gene promoter regions in dogs. PLoS Genet 9:e1003984. doi:10.1371/journal.pgen.1003984

Baudat F, Buard J, Grey C, Fledel-Alon A, Ober C, Przeworski M, Coop G, de Massy B. 2010. PRDM9 is a major determinant of meiotic recombination hotspots in humans and mice. Science 327:836–840. doi:10.1126/science.1183439

Bellacosa A, Drohat AC. 2015. Role of base excision repair in maintaining the genetic and epigenetic integrity of CpG sites. DNA Repair (Amst) 32:33–42. doi:10.1016/j.dnarep.2015.04.011

Bernardi G. 2000. Isochores and the evolutionary genomics of vertebrates. Gene 241:3–17. doi:10.1016/S0378-1119(99)00485-0

Bill CA, Duran WA, Miselis NR, Nickoloff JA. 1998. Efficient repair of all types of single-base mismatches in recombination intermediates in Chinese hamster ovary cells. Competition between long-patch and G-T glycosylase-mediated repair of G-T mismatches. Genetics 149:1935–1943.

Brick K, Smagulova F, Khil P, Camerini-Otero RD, Petukhova GV. 2012. Genetic recombination is directed away from functional genomic elements in mice. Nature 485:642–645. doi:10.1038/nature11089

Carninci P, Sandelin A, Lenhard B, Katayama S, Shimokawa K, Ponjavic J, Semple CAM, Taylor MS, Engström PG, Frith MC, Forrest ARR, Alkema WB, Tan SL, Plessy C, Kodzius R, Ravasi T, Kasukawa T, Fukuda S, Kanamori-Katayama M, Kitazume Y, Kawaji H, Kai C, Nakamura M, Konno H, Nakano K, Mottagui-Tabar S, Arner P, Chesi A, Gustincich S, Persichetti F, Suzuki H, Grimmond SM, Wells CA, Orlando V, Wahlestedt C, Liu ET, Harbers M, Kawai J, Bajic VB, Hume DA, Hayashizaki Y. 2006. Genome-wide analysis of mammalian promoter architecture and evolution. Nat Genet 38:626–635. doi:10.1038/ng1789

Cenik C, Chua HN, Zhang H, Tarnawsky S, Akef A, Derti A, Tasan M, Moore MJ, Palazzo AF, Roth FP. 2011. Genome analysis reveals interplay between 5’UTR introns and nuclear mRNA export for secretory and mitochondrial genes. PLoS Genetics 7:e1001366. doi:10.1371/journal.pgen.1001366

dos Reis M, Wernisch L. 2009. Estimating Translational Selection in Eukaryotic Genomes. Molecular Biology and Evolution 26:451–461. doi:10.1093/molbev/msn272

Duret L, Galtier N. 2009. Biased gene conversion and the evolution of mammalian genomic landscapes. Annu Rev Genomics Hum Genet 10:285–311. doi:10.1146/annurev-genom-082908-150001

Fenouil R, Cauchy P, Koch F, Descostes N, Cabeza JZ, Innocenti C, Ferrier P, Spicuglia S, Gut M, Gut I, Andrau J-C. 2012. CpG islands and GC content dictate nucleosome depletion in a transcription-independent manner at mammalian promoters. Genome Res 22:2399– 2408. doi:10.1101/gr.138776.112

Figuet E, Ballenghien M, Romiguier J, Galtier N. 2015. Biased Gene Conversion and GC-Content Evolution in the Coding Sequences of Reptiles and Vertebrates. Genome Biology and Evolution 7:240–250. doi:10.1093/gbe/evu277

Galtier N, Roux C, Rousselle M, Romiguier J, Figuet E, Glémin S, Bierne N, Duret L. 2018. Codon Usage Bias in Animals: Disentangling the Effects of Natural Selection, Effective Population Size, and GC-Biased Gene Conversion. Mol Biol Evol 35:1092–1103. doi:10.1093/molbev/msy015

Gould SJ, Lewontin RC. 1979. The spandrels of San Marco and the Panglossian paradigm: a critique of the adaptationist programme. *Proc R Soc Lond, B*, Biol Sci 205:581–598.

Haerty W, Ponting CP. 2015. Unexpected selection to retain high GC content and splicing enhancers within exons of multiexonic lncRNA loci. RNA 21:320–332. doi:10.1261/rna.047324.114

Hoge C, Manuel M de, Mahgoub M, Okami N, Fuller Z, Banerjee S, Baker Z, McNulty M, Andolfatto P, Macfarlan TS, Schumer M, Tzika AC, Przeworski M. 2023. Patterns of recombination in snakes reveal a tug of war between PRDM9 and promoter-like features. doi:10.1101/2023.07.11.548536

Huang Y, Gattoni R, Stévenin J, Steitz JA. 2003. SR splicing factors serve as adapter proteins for TAP-dependent mRNA export. Mol Cell 11:837–843.

Huang Y, Steitz JA. 2001. Splicing factors SRp20 and 9G8 promote the nucleocytoplasmic export of mRNA. Mol Cell 7:899–905.

International HapMap Consortium, Frazer KA, Ballinger DG, Cox DR, Hinds DA, Stuve LL, Gibbs RA, Belmont JW, Boudreau A, Hardenbol P, Leal SM, Pasternak S, Wheeler DA, Willis TD, Yu F, Yang H, Zeng C, Gao Y, Hu H, Hu W, Li C, Lin W, Liu S, Pan H, Tang X, Wang J, Wang W, Yu J, Zhang B, Zhang Q, Zhao Hongbin, Zhao Hui, Zhou J, Gabriel SB, Barry R, Blumenstiel B, Camargo A, Defelice M, Faggart M, Goyette M, Gupta S, Moore J, Nguyen H, Onofrio RC, Parkin M, Roy J, Stahl E, Winchester E, Ziaugra L, Altshuler D, Shen Yan, Yao Z, Huang W, Chu X, He Y, Jin L, Liu Y, Shen Yayun, Sun W, Wang Haifeng, Wang Yi, Wang Ying, Xiong X, Xu L, Waye MMY, Tsui SKW, Xue H, Wong JT-F, Galver LM, Fan J-B, Gunderson K, Murray SS, Oliphant AR, Chee MS, Montpetit A, Chagnon F, Ferretti V, Leboeuf M, Olivier J-F, Phillips MS, Roumy S, Sallée C, Verner A, Hudson TJ, Kwok P-Y, Cai D, Koboldt DC, Miller RD, Pawlikowska L, Taillon-Miller P, Xiao M, Tsui L-C, Mak W, Song YQ, Tam PKH, Nakamura Y, Kawaguchi T, Kitamoto T, Morizono T, Nagashima A, Ohnishi Y, Sekine A, Tanaka T, Tsunoda T, Deloukas P, Bird CP, Delgado M, Dermitzakis ET, Gwilliam R, Hunt S, Morrison J, Powell D, Stranger BE, Whittaker P, Bentley DR, Daly MJ, de Bakker PIW, Barrett J, Chretien YR, Maller J, McCarroll S, Patterson N, Pe’er I, Price A, Purcell S, Richter DJ, Sabeti P, Saxena R, Schaffner SF, Sham PC, Varilly P, Altshuler D, Stein LD, Krishnan L, Smith AV, Tello-Ruiz MK, Thorisson GA, Chakravarti A, Chen PE, Cutler DJ, Kashuk CS, Lin S, Abecasis GR, Guan W, Li Y, Munro HM, Qin ZS, Thomas DJ, McVean G, Auton A, Bottolo L, Cardin N, Eyheramendy S, Freeman C, Marchini J, Myers S, Spencer C, Stephens M, Donnelly P, Cardon LR, Clarke G, Evans DM, Morris AP, Weir BS, Tsunoda T, Mullikin JC, Sherry ST, Feolo M, Skol A, Zhang H, Zeng C, Zhao Hui, Matsuda I, Fukushima Y, Macer DR, Suda E, Rotimi CN, Adebamowo CA, Ajayi I, Aniagwu T, Marshall PA, Nkwodimmah C, Royal CDM, Leppert MF, Dixon M, Peiffer A, Qiu R, Kent A, Kato K, Niikawa N, Adewole IF, Knoppers BM, Foster MW, Clayton EW, Watkin J, Gibbs RA, Belmont JW, Muzny D, Nazareth L, Sodergren E, Weinstock GM, Wheeler DA, Yakub I, Gabriel SB, Onofrio RC, Richter DJ, Ziaugra L, Birren BW, Daly MJ, Altshuler D, Wilson RK, Fulton LL, Rogers J, Burton J, Carter NP, Clee CM, Griffiths M, Jones MC, McLay K, Plumb RW, Ross MT, Sims SK, Willey DL, Chen Z, Han H, Kang L, Godbout M, Wallenburg JC, L’Archevêque P, Bellemare G, Saeki K, Wang Hongguang, An D, Fu H, Li Q, Wang Z, Wang R, Holden AL, Brooks LD, McEwen JE, Guyer MS, Wang VO, Peterson JL, Shi M, Spiegel J, Sung LM, Zacharia LF, Collins FS, Kennedy K, Jamieson R, Stewart J. 2007. A second generation human haplotype map of over 3.1 million SNPs. Nature 449:851–861. doi:10.1038/nature06258

Jónsson H, Sulem P, Kehr B, Kristmundsdottir S, Zink F, Hjartarson E, Hardarson MT, Hjorleifsson KE, Eggertsson HP, Gudjonsson SA, Ward LD, Arnadottir GA, Helgason EA, Helgason H, Gylfason A, Jonasdottir Adalbjorg, Jonasdottir Aslaug, Rafnar T, Frigge M, Stacey SN, Th. Magnusson O, Thorsteinsdottir U, Masson G, Kong A, Halldorsson BV, Helgason A, Gudbjartsson DF, Stefansson K. 2017. Parental influence on human germline de novo mutations in 1,548 trios from Iceland. Nature 549:519–522. doi:10.1038/nature24018

Joseph J, Prentout D, Laverré A, Tricou T, Duret L. 2023. High prevalence of Prdm9-independent recombination hotspots in placental mammals (preprint). Genetics. doi:10.1101/2023.11.17.567540

Kaessmann H, Vinckenbosch N, Long M. 2009. RNA-based gene duplication: mechanistic and evolutionary insights. Nat Rev Genet 10:19–31. doi:10.1038/nrg2487

Kalari KR, Casavant M, Bair TB, Keen HL, Comeron JM, Casavant TL, Scheetz TE. 2006. First exons and introns--a survey of GC content and gene structure in the human genome. In Silico Biol (Gedrukt) 6:237–242.

Koonin EV. 2016. Splendor and misery of adaptation, or the importance of neutral null for understanding evolution. BMC Biol 14:114. doi:10.1186/s12915-016-0338-2

Lei H, Dias AP, Reed R. 2011. Export and stability of naturally intronless mRNAs require specific coding region sequences and the TREX mRNA export complex. Proc Natl Acad Sci USA 108:17985–17990. doi:10.1073/pnas.1113076108

Lei H, Zhai B, Yin S, Gygi S, Reed R. 2013. Evidence that a consensus element found in naturally intronless mRNAs promotes mRNA export. Nucleic Acids Res 41:2517–2525. doi:10.1093/nar/gks1314

Lesecque Y, Glémin S, Lartillot N, Mouchiroud D, Duret L. 2014. The Red Queen Model of Recombination Hotspots Evolution in the Light of Archaic and Modern Human Genomes. PLoS Genet 10:e1004790. doi:10.1371/journal.pgen.1004790

Louie E, Ott J, Majewski J. 2003. Nucleotide frequency variation across human genes. Genome Res 13:2594–2601. doi:10.1101/gr.1317703

Lynch M. 2007. The frailty of adaptive hypotheses for the origins of organismal complexity. Proceedings of the National Academy of Sciences 104:8597–8604. doi:10.1073/pnas.0702207104

Masuda S, Das R, Cheng H, Hurt E, Dorman N, Reed R. 2005. Recruitment of the human TREX complex to mRNA during splicing. Genes Dev 19:1512–1517. doi:10.1101/gad.1302205

Mihola O, Pratto F, Brick K, Linhartova E, Kobets T, Flachs P, Baker CL, Sedlacek R, Paigen K, Petkov PM, Camerini-Otero RD, Trachtulec Z. 2019. Histone methyltransferase PRDM9 is not essential for meiosis in male mice. Genome Res 29:1078–1086. doi:10.1101/gr.244426.118

Mordstein C, Savisaar R, Young RS, Bazile J, Talmane L, Luft J, Liss M, Taylor MS, Hurst LD, Kudla G. 2020. Codon Usage and Splicing Jointly Influence mRNA Localization. Cell Systems 10:351–362.e8. doi:10.1016/j.cels.2020.03.001

Myers S, Bowden R, Tumian A, Bontrop RE, Freeman C, MacFie TS, McVean G, Donnelly P. 2010. Drive Against Hotspot Motifs in Primates Implicates the PRDM9 gene in Meiotic Recombination. Science 327:10.1126/science.1182363. doi:10.1126/science.1182363

Paiano J, Wu W, Yamada S, Sciascia N, Callen E, Paola Cotrim A, Deshpande RA, Maman Y, Day A, Paull TT, Nussenzweig A. 2020. ATM and PRDM9 regulate SPO11-bound recombination intermediates during meiosis. Nat Commun 11:857. doi:10.1038/s41467-020-14654-w

Paigen K, Petkov PM. 2018. PRDM9 and its role in genetic recombination. Trends Genet 34:291–300. doi:10.1016/j.tig.2017.12.017

Palazzo A, Mahadevan K, Tarnawsky S. 2013. ALREX-elements and introns: two identity elements that promote mRNA nuclear export. WIREs RNA 4:523–533. doi:10.1002/wrna.1176

Palazzo AF, Akef A. 2012. Nuclear export as a key arbiter of “mRNA identity” in eukaryotes. Biochim Biophys Acta 1819:566–577. doi:10.1016/j.bbagrm.2011.12.012

Palazzo AF, Kang YM. 2021. GC-content biases in protein-coding genes act as an “mRNA identity” feature for nuclear export. BioEssays 43:2000197. doi:10.1002/bies.202000197

Palazzo AF, Kejiou NS. 2022. Non-Darwinian Molecular Biology. Frontiers in Genetics 13:831068.

Palazzo AF, Qiu Y, Kang YM. 2024. mRNA nuclear export: how mRNA identity features distinguish functional RNAs from junk transcripts. RNA Biology 21:1–12. doi:10.1080/15476286.2023.2293339

Palazzo AF, Springer M, Shibata Y, Lee C-S, Dias AP, Rapoport TA. 2007. The signal sequence coding region promotes nuclear export of mRNA. PLoS Biol 5:e322. doi:10.1371/journal.pbio.0050322

Polak P, Arndt PF. 2008. Transcription induces strand-specific mutations at the 5′ end of human genes. Genome Res 18:1216–1223. doi:10.1101/gr.076570.108

Pouyet F, Gilbert KJ. 2021. Towards an improved understanding of molecular evolution: the relative roles of selection, drift, and everything in between. Peer Community Journal 1. doi:10.24072/pcjournal.16

Pouyet F, Mouchiroud D, Duret L, Sémon M. 2017. Recombination, meiotic expression and human codon usage. eLife 6:e27344. doi:10.7554/eLife.27344

Roca X, Sachidanandam R, Krainer AR. 2005. Determinants of the inherent strength of human 5′ splice sites. RNA 11:683–698. doi:10.1261/rna.2040605

Schield DR, Pasquesi GIM, Perry BW, Adams RH, Nikolakis ZL, Westfall AK, Orton RW, Meik JM, Mackessy SP, Castoe TA. 2020. Snake Recombination Landscapes Are Concentrated in Functional Regions despite PRDM9. Molecular Biology and Evolution 37:1272–1294. doi:10.1093/molbev/msaa003

Singhal S, Leffler EM, Sannareddy K, Turner I, Venn O, Hooper DM, Strand AI, Li Q, Raney B, Balakrishnan CN, Griffith SC, McVean G, Przeworski M. 2015. Stable recombination hotspots in birds. Science 350:928–932. doi:10.1126/science.aad0843

Smagulova F, Gregoretti IV, Brick K, Khil P, Camerini-Otero RD, Petukhova GV. 2011. Genome-wide analysis reveals novel molecular features of mouse recombination hotspots. Nature 472:375–378. doi:10.1038/nature09869

Tamarkin-Ben-Harush A, Vasseur J-J, Debart F, Ulitsky I, Dikstein R. 2017. Cap-proximal nucleotides via differential eIF4E binding and alternative promoter usage mediate translational response to energy stress. Elife 6:e21907. doi:10.7554/eLife.21907

Tarnawsky SP, Palazzo AF. 2012. Positional requirements for the stimulation of mRNA nuclear export by ALREX-promoting elements. Mol Biosyst 8:2527–2530. doi:10.1039/c2mb25016k

Thomas A, Rehfeld F, Zhang H, Chang T-C, Goodarzi M, Gillet F, Mendell JT. 2022. RBM33 directs the nuclear export of transcripts containing GC-rich elements. Genes Dev 36:550–565. doi:10.1101/gad.349456.122

Walsh CP, Xu GL. 2006. Cytosine methylation and DNA repair. Curr Top Microbiol Immunol 301:283–315. doi:10.1007/3-540-31390-7_11

Williams AL, Genovese G, Dyer T, Altemose N, Truax K, Jun G, Patterson N, Myers SR, Curran JE, Duggirala R, Blangero J, Reich D, Przeworski M, T2D-GENES Consortium. 2015. Non-crossover gene conversions show strong GC bias and unexpected clustering in humans. Elife 4. doi:10.7554/eLife.04637

Xia X, Xie Z, Li W-H. 2003. Effects of GC content and mutational pressure on the lengths of exons and coding sequences. J Mol Evol 56:362–370. doi:10.1007/s00239-002-2406-1

Xie Y, Gao S, Zhang K, Bhat P, Clarke BP, Batten K, Mei M, Gazzara M, Shay JW, Lynch KW, Angelos AE, Hill PS, Ivey AL, Fontoura BMA, Ren Y. 2023. Structural basis for high-order complex of SARNP and DDX39B to facilitate mRNP assembly. Cell Rep 42:112988. doi:10.1016/j.celrep.2023.112988

Zhang L, Kasif S, Cantor CR, Broude NE. 2004. GC/AT-content spikes as genomic punctuation marks. Proc Natl Acad Sci USA 101:16855–16860. doi:10.1073/pnas.0407821101

Zhu L, Zhang Y, Zhang W, Yang S, Chen J-Q, Tian D. 2009. Patterns of exon-intron architecture variation of genes in eukaryotic genomes. BMC Genomics 10:47. doi:10.1186/1471-2164-10-47

Zuckerman B, Ron M, Mikl M, Segal E, Ulitsky I. 2020. Gene Architecture and Sequence Composition Underpin Selective Dependency of Nuclear Export of Long RNAs on NXF1 and the TREX Complex. Molecular Cell 79:251–267.e6. doi:10.1016/j.molcel.2020.05.013

